# Elucidating the mechanisms of action and evolutionary relationships of phage anti-defence proteins

**DOI:** 10.1101/2025.06.06.658234

**Authors:** Khalimat Murtazalieva, Evangelos Karatzas, Alessio Yang, Federico Corona, Jiawei Wang, Athanasios Typas, Robert D. Finn

**Author notes:** These authors contributed equally to this work (Alessio Yang, Federico Corona). Correspondence (K.M.), (R.D.F.).

## Abstract

Phages and bacteria are locked in a molecular arms race, with phage anti-defence proteins (ADPs) enabling them to evade bacterial immune systems. To streamline access to information on ADPs, we developed the Encyclopaedia of Viral Anti-DefencE Systems (EVADES), an online resource containing sequences, structures, protein family annotations, and mechanisms of action (MoA) for 268 ADPs. Through computational structural analysis we predicted MoAs for 21 uncharacterised ADPs. We demonstrate the utility of EVADES by exploring different characteristics of ADPs: (i) DNA mimic ADPs exhibit broad-spectrum activity against bacterial defence systems; (ii) protein sharing across defence systems enables multi-defence inhibitory activity of ADPs; and (iii) eukaryotic dsDNA viruses encode ADP homologs, suggesting conserved immune evasion strategies across domains of life. These results highlight the broader relevance of phage ADPs in understanding interactions between prokaryotic or eukaryotic viruses and their hosts. EVADES is freely accessible at https://www.ebi.ac.uk/finn-srv/evades.

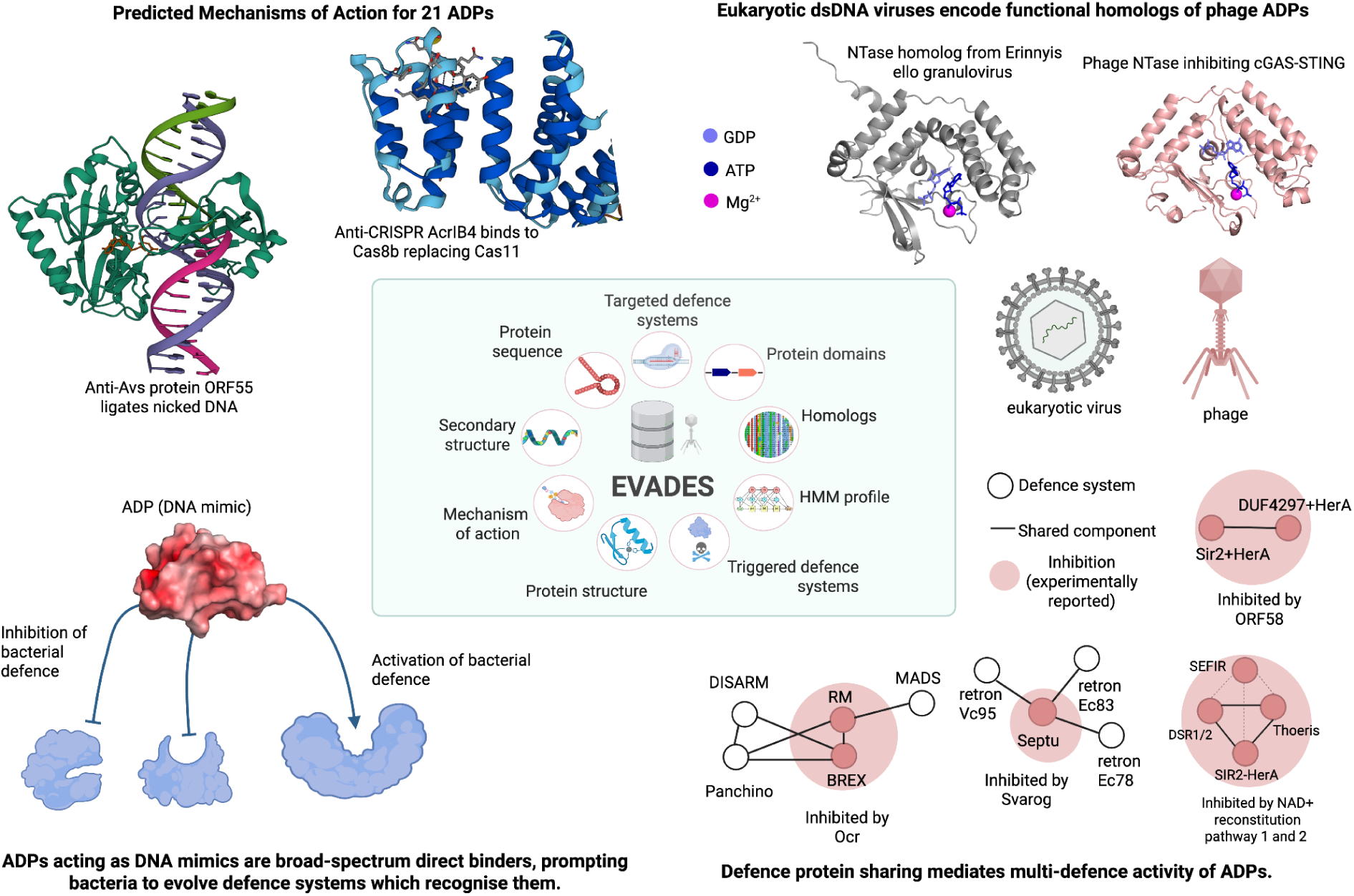

- The EVADES database aggregates knowledge on phage anti-defence proteins (ADPs), facilitating hypothesis generation.
- Mechanisms of action are proposed for 21 uncharacterised ADPs.
- ADPs that function as DNA mimics can block diverse bacterial defence systems.
- Homology of targets of ADPs underpin their broad counter-defence capacities and enable prediction of further targets.
- Eukaryotic double-stranded DNA viruses employ functional homologs of ADPs to evade host immune responses.

## INTRODUCTION

Biological arms races—the adaptation between competing sets of co-evolving genes or organisms—are ubiquitous in nature. One such example is the perpetual arms race between bacteria and their viruses, bacteriophages (hereafter termed phages), which has driven the evolution of myriads of defence systems in bacteria that help them protect against phage infections^1^. Currently, known bacterial defence systems employ a wide range of molecular mechanisms to prevent phage replication. These include degradation of phage nucleic acids, interference with or blocking essential molecular processes for the phage, or mechanisms that involve targeting the host cell before the phage manages to replicate (collectively called abortive infection)^1,2^. In response, phages have developed an array of anti-defence systems aimed at circumventing or neutralising these bacterial defences, thereby ensuring successful infection and replication^3^.

Since the discovery of the first anti-defence proteins (ADPs) against restriction-modification (RM) enzymes in the 1970s^4,5^, and the explosion of anti-CRISPR mechanism discovery in the 2010’s^6^, the number of characterised systems has grown significantly^7^, particularly over the past decade. This underscores the need for a resource that systematically classifies and describes ADPs to enhance our understanding of their mechanisms and evolutionary diversity. In this study, we compiled a comprehensive collection of ADPs, including their sequences, experimental or predicted secondary and tertiary structures, functional domains, and mechanisms of action (MoA). However, many entries have not been functionally characterised in terms of MoA, questioning whether they directly target defence systems or act via indirect mechanisms. Recognising this uncertainty, we incorporated Evidence Code Ontology (ECO) labels^8^ to indicate the type of evidence supporting each entry, thereby improving interpretability and confidence in the data. Complementing this, we conducted computational analyses and proposed putative MoAs for 21 previously uncharacterised ADPs. Multitarget activity is an emerging theme among phage ADPs. Striking examples of broad acting ADPs are phage kinases that counteract defence systems via promiscuous phosphorylation. The T7 protein kinase exhibits anti-defense activity against many nucleic-acid binding defense systems (e.g. Retron-Eco9 and DarTG1) of *E. coli*^9^, and so does its homologous *Salmonella* phage JSS protein kinase (Dnd, CRISPR‒Cas, QatABCD, SIR2+HerA and DUF4297+HerA systems)^10^. ADPs can also act on conserved components or functions shared by different bacterial defence systems. Well-characterised examples include: (i) Thoeris anti-defence proteins 1 and 2 (Tad1/2)^11–13^, which inhibit both Thoeris and CBASS systems by sequestering cyclic nucleotides; (ii) Ocr, which binds to PglX^14^ of the Bacteriophage Exclusion (BREX) system—and homologous HsdM methyltransferase in RM defence; (iii) S-adenosyl-methionine (SAM) lyase^15^, which cleaves SAM, a molecule utilised by RM and BREX defences; and (iv) NAD+ reconstitution pathways 1 and 2 (NARP1/2), which evade many defence systems that deplete NAD+, including Thoeris, DSR1, DSR2, SIR2-HerA, and SEFIR^16^. Lastly, there are also ADPs that have poorly understood multitarget activity. ORF58^17^, with an uncharacterised MoA, evades both Sir2+HerA and DUF4297+HerA defences, which share structural similarity^18^. We hypothesise that multipurpose ADPs may be widespread, as bacterial defence systems have evolved through protein domain shuffling across different defence systems^19^.

Many ADPs are sensed by other defense systems as triggers^20–24^. These cascade relationships are likely a general wiring principle of bacterial defence systems, which have learned to detect anti-defence activities^25^. Activation of a bacterial defense system by a phage ADP may occur through direct interaction with the defence components. For instance, the PARIS defence senses the DNA mimic Ocr from *Escherichia* phage T7 through binding^26^ and various retrons are activated by DNA methylation (Dam or Dcm), which is used by phages to circumvent RM systems^23,27^. Bacterial defences may also recognise phage ADPs indirectly. For example, it has been proposed that the retrons Se72 and Ec48 “guard” the RecBCD system in *E. coli*, detecting the inhibition of the RecBCD complex—for instance, by phage *λ* Gam protein or the *T7* Gp5.9 protein—and triggering abortive infection^28^. The Panoptes defence synthesises decoy signalling nucleotides, which are sensed by its effector, OptE, preventing membrane disruption. The phage sponge protein Acb2 activates the Panoptes defence by sequestering these signalling nucleotides and thereby triggering the OptE effector protein^29^. To better understand the properties of ADPs that trigger bacterial defence systems, we examined their features to identify commonalities.

Bacterial defense systems seem to also be ancestral to different components of eukaryotic immunity^30,31^. For instance, nucleotide-binding oligomerization domain–like receptors (NLRs) of the STAND superfamily (receptors in eukaryotes involved in innate immunity) are homologous to the bacterial Avs-defence system, which recognises conserved viral proteins^32^. Evolutionary links between phages and eukaryotic dsDNA viruses have been also extensively described^33–35^. For example, it has been hypothesised that polintoviruses were the first group of eukaryotic dsDNA viruses that evolved from bacteriophages, and that they subsequently gave rise to most large DNA viruses of eukaryotes^33^. This shared evolutionary history, together with the structural and functional homology between the phage anti-CBASS protein 1 (Acb1) and the poxvirus cGAMP phosphodiesterase (inhibits the animal cGLR response)^36^, implies that ADPs are conserved between bacterial and eukaryotic viruses. Consistently, phage-encoded Thoeris anti-defence protein 4 (Tad4) and anti-CBASS protein 3 (Acb3) can inhibit related innate immune systems in plants, animals and humans^37^, suggesting that retaining phage-evolved ADPs could be beneficial for eukaryotic viruses. Hence, we decided to investigate further the evolutionary and functional relationships of ADPs that counteract defence systems shared between bacteria and eukaryotes.

In this study, we systematically survey the literature for phage ADPs, investigate their functions, targets and potential to act as triggers for other defense systems using bioinformatics. We also extend the systematic comparisons to systems evading eukaryotic immunity. Since ADP discovery is still at its infancy, we use phage genomic information (diversity islands containing anti-defence genes) and AlphaFold 3^38^ to predict ultra-high confidence direct binders among uncharacterised phage proteins to effectors of bacterial defence systems. By uniting structural, evolutionary, and functional insights, our work provides a foundation for advancing the understanding of phage ADPs and their roles in viral–host interplay across domains of life.

## RESULTS

### Diversity of phage ADPs

We have established the Encyclopaedia of Viral Anti-DefencE Systems (EVADES), which catalogues 268 ADPs that form components of 247 anti-defence systems^9–13,15–18,21,23,24,32,37,39–146^, some of which are multicomponent systems (see Methods – Data Collection). Collectively, they target 45 families of bacterial defences. The EVADES database encapsulates a significantly broader diversity of phage ADPs compared to other resources (Figure 1A, Supplementary Table 1). For example, AntiDefenseFinder catalogues 176 ADPs across 156 systems^147^. The dbAPIS database describes 80 experimentally validated ADPs, along with their remote homologs^148^. In contrast, resources such as Anti-CRISPRdb^149,150^, and AcrHub^151^ focus exclusively on anti-CRISPR (Acr) proteins, with AcrHub containing 76 such proteins.

**Figure 1.**
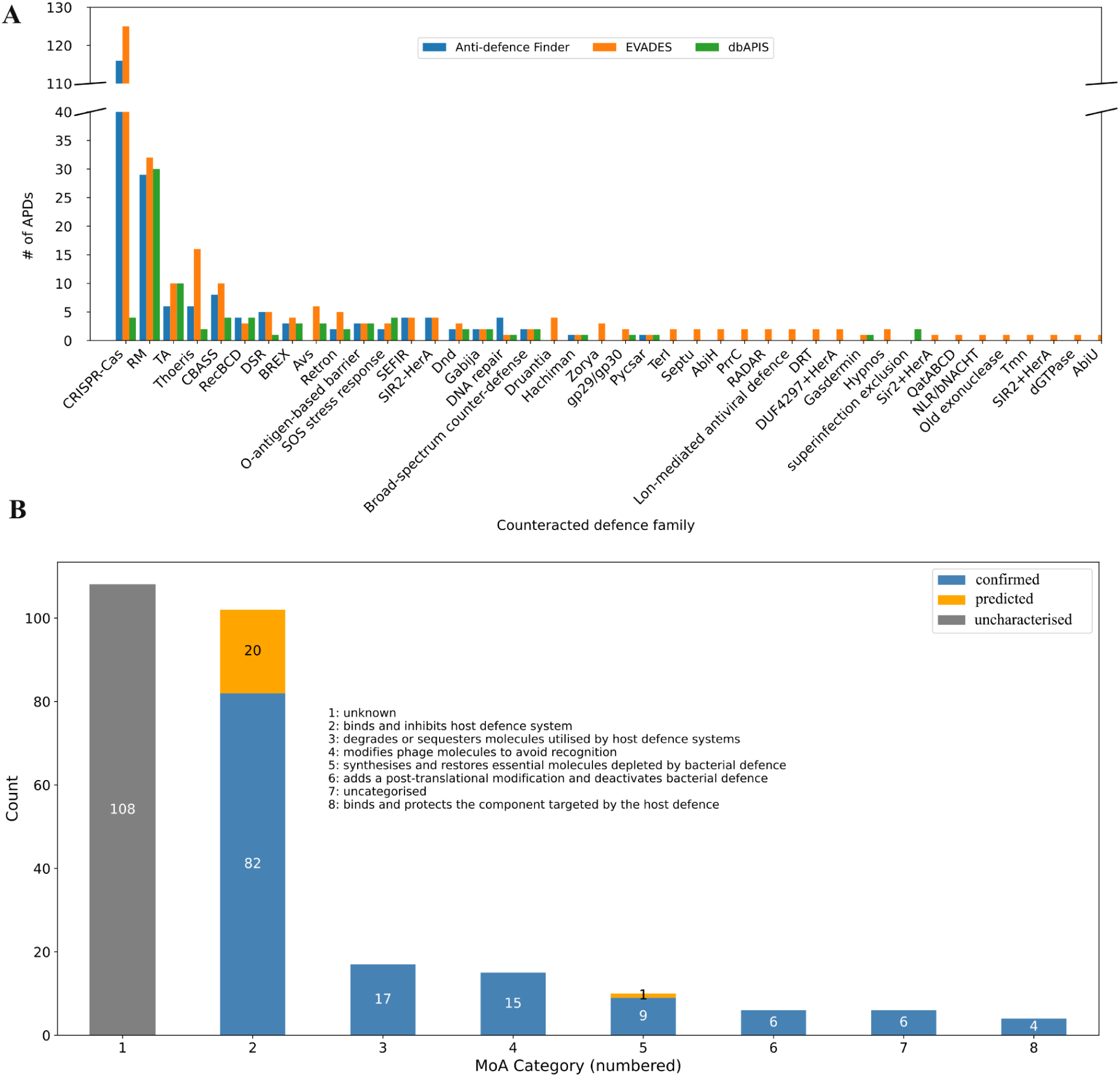
Overview of the database. **(A)** Distribution of anti-defence proteins (ADPs) in AntiDefenseFinder (blue), EVADES (orange) and dbAPIS (green) by the family of targeted defence systems. The Y-axis was truncated due to disproportionately high counts of anti-CRISPR proteins compared to other defence systems. **(B)** Distribution of ADPs in EVADES. Bars represent the number of proteins assigned to each MoA category (1–8), split to experimentally confirmed MoAs (blue) and newly predicted ones from this study (orange). The total count for each category is displayed on the corresponding bar.

Beyond its broad range of ADPs entries, EVADES provides detailed MoAs, ECO labels, protein sequences, experimental and predicted structures, profile-hidden Markov models (HMMs), information on protein family membership and homologs in eukaryotic dsDNA viruses, and additional data—including the potential to trigger other defence systems. Only dbAPIS^148^ and AcrHub^151^ provide comprehensive information, but are restricted in their scopes.

We classified known ADPs into categories based on their MoAs (Figure 1B and Supplementary Figure 1): (i) ADPs with an unknown MoA (n = 108); (ii) ADPs that bind to bacterial defence components (n = 102), including 20 new AlphaFold 3 predictions from this study; (iii) ADPs that degrade or sequester signalling molecules/cofactors utilised by bacterial defences (n =17); (iv) ADPs that modify phage components to evade recognition (n = 15); (v) ADPs that synthesise or restore essential molecules depleted by bacterial defences, including one predicted in this study (n=10); (vi) ADPs that add post-translational modifications to bacterial defence systems (n = 6); (vii) uncategorised ADPs (n = 6); and (viii) proteins that bind and protect components targeted by host defences (n = 4). While 70 out of 136 ADPs with experimentally characterised MoAs have known structures in the Protein Data Bank (PDB)^152^, the vast majority of uncharacterised ADPs (119 out of 129) lack structural data. To address this and gain insights into potential MoAs, we predicted protein structures for all ADPs lacking a known structure and performed structural similarity searches (see methods).

We also compared the sequences of ADPs to infer evolutionary relationships between them and found many pairs with high similarity at the sequence level (Supplementary Figure 2A). Interestingly, some of these pairs comprised proteins that counteract different defence families. For instance, Tad2^12^ and AcrIIA7^54^ shared high sequence and structure-based similarity (Supplementary Figure 2B-C, p-value = 2.55 × 10⁻⁸). Tad2 sequesters gcADPR^12^, a signalling molecule utilised by the Thoeris defence system. AcrIIA7 has been reported as type II-A anti-CRISPR protein with unknown MoA^54^. As type II CRISPR–Cas systems are not known to utilise cyclic signalling molecules, this similarity was unexpected. Although AcrIIA7 contains a variable central fragment that aligns poorly with Tad2, the rest of the sequence, including conserved residues, align well (Supplementary Figure 2C). Moreover, AcrIIA7 encodes residues similar to those at Tad2’s active site. Based on these results, we concluded that AcrIIA7 is highly likely to sequester gcADPR, similar to Tad2 (Supplementary Figure 2B-C). During the course of this work, this functional prediction was validated by an experimental study^11^; however, the authors were unable to confirm the anti-CRISPR activity previously reported in a functional metagenomic screen^54^. Nevertheless, this example illustrates how structural computational biology approaches can elucidate molecular function based on similarity with other proteins. It also highlights the importance of including information on the nature of evidence supporting the anti-defence activity of database entries.

### Inferring multi-defence activity of ADPs from defence protein-sharing network

Multi-defence activity of ADPs can be mediated by exploiting similar protein components of different defence systems. To explore this phenomenon further, we constructed a network based on structural similarity among bacterial defence proteins. We focused on defence proteins that are either directly targeted by ADPs or are presumed to be targeted via uncharacterised mechanisms of action, and expanded it by including other defence proteins that shared significant similarity (Supplementary Table 2). We then added ADPs as triangular nodes to indicate inhibitory interactions. The presence of shared components among defence systems suggests that some ADPs may also inhibit other systems that exhibit similarity to their known targets (Supplementary Figure 3A). For instance, it has been shown that Ocr binds to PglX, a component of the BREX defence system. PglX shares high structural similarity with gp28 from the Panchino defence, drmMI from Defense Island System Associated with Restriction-Modification (DISARM), and HsdM from the type I RM system. While inhibition of the type I RM system by Ocr has been demonstrated—specifically via binding between the HsdM and HsdR subunits— we hypothesise that Ocr may also inhibit DISARM and Panchino defence systems by binding to drmMI and gp28, respectively. This hypothesis is supported by AlphaFold 3 predictions of complexes between Ocr and drmMI or gp28. While these predictions are of a moderate confidence (0.6 > ipTM score < 0.8), the binding interface of Ocr was similar in the two complexes (Supplementary Figure 3B).

### 20 ADPs with uncharacterised MoA are likely to act as direct binders

Since direct binders represent the most frequent characterised MoA category, we hypothesised that some of the ADPs with uncharacterised function may act via the same mechanism. To investigate this, we co-folded ADPs with uncharacterised MoAs alongside the proteins from the targeted defence systems (Supplementary Table 3). This analysis predicted 13 high-confidence interactions and 10 moderate-confidence interactions between ADPs and their corresponding defence systems (Supplementary Table 4).

While examining the results, we found that the complex between AcrIIA13 and *Staphylococcus aureus* Cas9 (*Sau*Cas9) had been confirmed by an almost identical structure (PDB accession: 7ENI). However, as this PDB entry lacked a corresponding publication, it had not been initially included in our database. Additionally, we identified a recent paper that experimentally confirmed the interaction between Cas5e and AcrIE3—predicted by AlphaFold 3, with an interface predicted template modelling (ipTM) score of 0.84 and predicted template modelling (pTM) score of 0.84—which had also not been included in our database^63^. We subsequently updated the MoA annotations for AcrIIA13 and AcrIE3, and added the other predictions, indicating that they are based on structural models.

We analysed our predictions alongside the original publications reporting ADPs to gain mechanistic insights and assess whether our predicted results align with available experimental data. For example, we predicted a high-confidence complex (ipTM = 0.92, pTM = 0.85) between AcrIC5 and *Pseudomonas aeruginosa* Cas8c (*Pa*Cas8c, NCBI GenPept accession: UEM35121.1). AcrIC5 has been reported to block *P. aeruginosa* CRISPR interference (CRISPRi), suggesting that it prevents target DNA binding^40^. Consistent with this, our prediction places AcrIC5 near the protospacer adjacent motif (PAM) binding site on PaCas8^153^. Notably, AcrIC5 is a highly acidic molecule (Figure 2A). The interaction between the PAM motif in the target DNA and the PAM-binding site on Cas proteins is essential for DNA binding. AcrIC5 may therefore displace the DNA and sterically block its binding. This prediction is consistent with the observation that AcrIC5 inhibits CRISPRi by preventing target DNA binding^40^. In contrast to AcrIC5, AcrIC3 was shown to be unable to inhibit CRISPRi in the same study^40^, suggesting that it acts downstream of DNA binding. Consistent with a downstream target (obstructing Cas3 recruitment or DNA cleavage), we predicted a high-confidence (ipTM = 0.91, pTM = 0.90) interaction between the Cas3 endonuclease (GenBank: UEM35119.1)—spanning all four of its domains—and AcrIC3 (Figure 2B).

**Figure 2.**
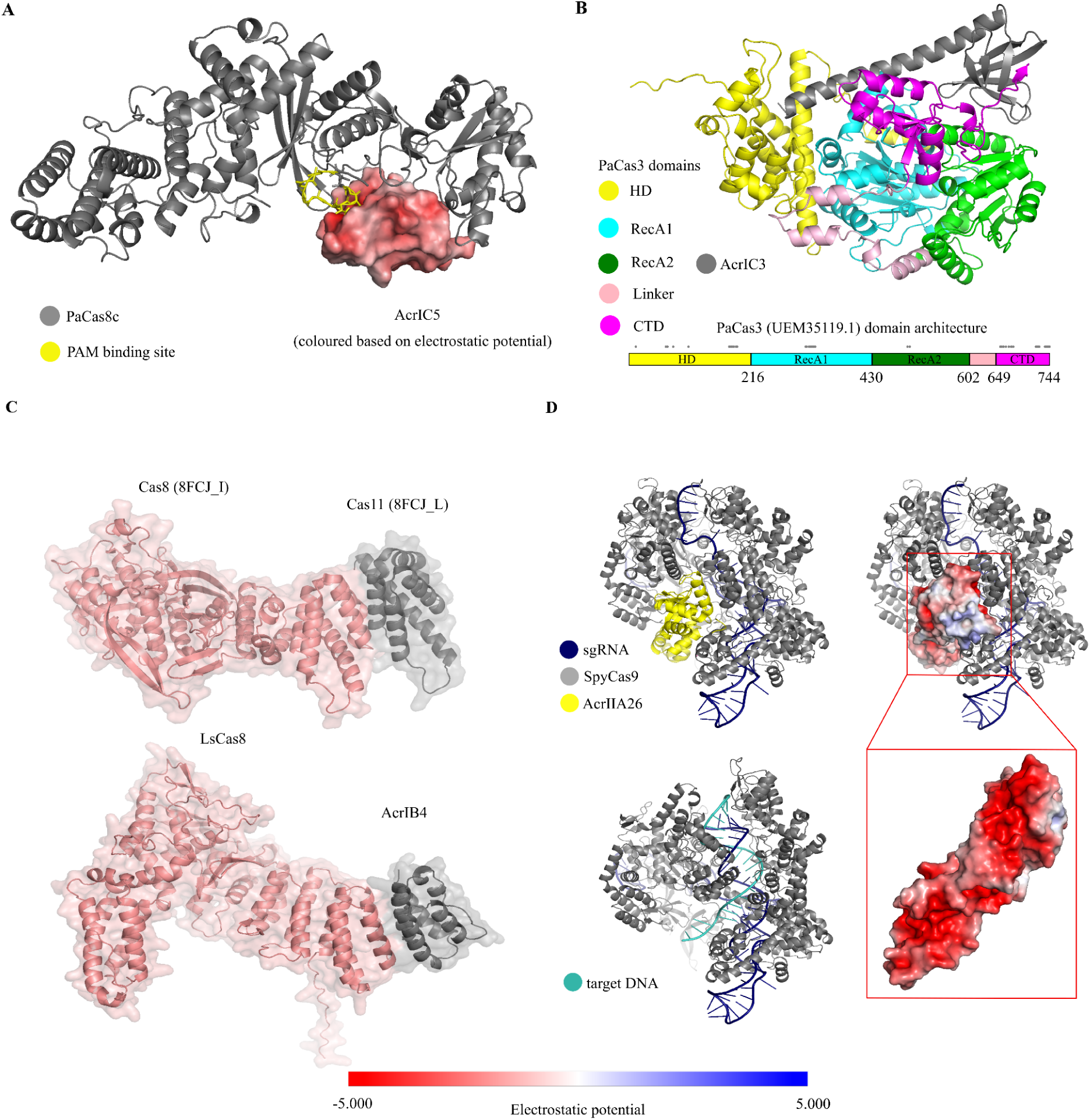
Putative mechanisms of action for ADPs based on structural model analysis. **(A)** AcrIC5 (surface rendered and coloured based on electrostatic potential – colour code is shown below the entire Figure) is a negatively charged protein that binds to Cas8c (grey, ribbon representations), occluding its protospacer adjacent motif (PAM)-binding site (yellow, side chains shown a sticks) and preventing Cas8c from interacting with the PAM site on the target DNA. **(B)** AcrIC3 (grey) binds to Cas3, with interactions spanning all four domains of Cas3: the HD nuclease (yellow), RecA1 (cyan), RecA2 (green), and C-terminal domain (CTD, magenta). Underneath the structure is a linear representation of Cas3 domain architecture, which is coloured as in the structure. Grey dots indicate the regions involved in interacting with AcrIC3. **(C)** Typically, the C-terminal domain of *Ls*Cas8b interacts with Cas11 (top), which stabilises Cas8 within the Cascade. AcrIB4 is likely to bind to Cas8b, replacing Cas11 in the Cascade. Cas8b from *Listeria seeligeri* (*Ls*Cas8b, coloured salmon pink) is predicted to bind AcrIB4 (grey) via its C-terminal domain (bottom). **(D)** AcrIIA26 (yellow, top-left) is predicted to form a high-confidence complex with *Streptococcus pyogenes* Cas9 (*Spy*Cas9, grey) and single guide RNA (cyan, sgRNA). The complex modelled between *Spy*Cas9, sgRNA, and target DNA (cyan, bottom-left) suggests that AcrIIA26 is likely to occlude the *Spy*Cas9’s DNA-binding site. In the complex between *Spy*Cas9, sgRNA, and AcrIIA26, the latter is coloured based on electrostatic potential (top-right). We rotated AcrIIA26 (135 degrees around y axis and then 55 angles around y axis, both in counterclockwise directions) to reveal the *Spy*Cas9 interaction interface (bottom-right).

AcrIB4 is a puzzling example of an anti-CRISPR protein, as it shares similarity to the C-terminal fragment of *Listeria seeligeri* Cas8b, raising the possibility that it may replace Cas8b during aberrant Cascade assembly. However, experiments have shown that AcrIB4 does not prevent the assembly of Cas8b or other core subunits (Cas7, Cas6, and Cas5) into the Cascade complex, and it has been reported to act upstream of target DNA binding^58^. The authors were unable to generate a functionally tagged version of AcrIB4 to directly assess its incorporation into the Cascade complex^58^. To understand the potential functional role of AcrIB4, we predicted a high-confidence interaction between AcrIB4 and the *Listeria seeligeri* Cas8b (ipTM = 0.91, pTM = 0.76), involving the C-terminal residues of Cas8b that mediate binding to Cas11 (Figure 2C). Cas11 is translated from an internal ribosome-binding site within the 3′ end of the *cas8b* gene and typically stabilises the non-target DNA strand during R-loop formation by positioning it above the crRNA–target DNA heteroduplex^154^. Based on this, we propose that AcrIB4 replaces Cas11 in the complex, thereby destabilising the R-loop formation.

AcrIIA26 has been characterised as a potent inhibitor of *Streptococcus pyogenes* Cas9 (*Spy*Cas9). It does not affect the formation of the Cas9–single guide RNA (sgRNA) complex but prevents this complex from binding to DNA, suggesting that it interferes at a downstream stage of the targeting process^41^. Consistent with this data, we predicted a high-confidence direct interaction (ipTM = 0.87, pTM = 0.79) between AcrIIA26 and *Spy*Cas9, involving the negatively charged surface of AcrIIA26 and residues in *Spy*Cas9, typically involved in target DNA binding (Figure 2D).

### Phage ADPs trigger bacterial defence systems

The mechanisms by which bacterial defence systems sense phage infection remain poorly understood. However, several studies have identified the molecular triggers of these systems^28,155^. Notably, Stokar-Avihail *et al.* conducted a systematic study aimed at uncovering phage determinants sensed by bacterial defences, including putative triggers of defence systems^28^. Their approach involved isolating phage mutants capable of escaping bacterial defences, sequencing them, and identifying mutations in either the protein-coding genes or in the intergenic regions adjacent to such genes. Such mutations were found in genes encoding structural proteins, components of the replication machinery, ADPs and proteins of unknown function. To extend this work, we compared entries in EVADES with those reported by Stokar-Avihail *et al.* to identify potential functions or relationships to newly discovered ADPs. We found that the putative Borvo trigger from phage SECphi18 is, in fact, the entry mga47 in EVADES, which has been reported as a DNA polymerase that is likely capable of replicating DNA ADP-ribosylated by the host DarTG system^130^. The putative ShosTA trigger (mutations resulting in the absence of the protein enable escape from the defense system) is identical to the T7 protein kinase, which inhibits the Retron-Eco9 and DarTG1 defence systems^9^, and 53.7% sequence identity over its full length with the *Salmonella* phage JSS1 protein kinase, which inhibits the Dnd, CRISPR‒Cas, QatABCD, SIR2+HerA, and DUF4297+HerA systems^10^. Additionally, among the putative triggers, the Avs type I trigger from phage SBSphiC caught our attention, as it was annotated as deoxyuridylate (dUMP) hydroxymethylase. It shared 97% sequence identity over its full length with the deoxyuridylate (dUMP) hydroxymethylase of *Bacillus* phage SP-10, as well as structural similarity to deoxycytidylate hydroxymethylase from phage T4 included in the database (FATCAT *p*-value = 4.88^e-12^). The dUMP hydroxymethylase of *Bacillus* phage SP-10 is involved in genome hypermodification and protection from restriction enzymes, similar to the deoxycytidylate hydroxymethylase from phage T4, suggesting that the Avs type I trigger may act as an ADP. We further searched for structural homologs of the type I Avs trigger in the PDB and found that it has extremely high similarity to cytidine monophosphate (CMP) hydroxymethylase MilA (Figure 3A, PDB: 5JNH; FATCAT 2.0 p-value =0.00^e+00^)^156^. However, there are indications that it may possess different substrate specificity from MilA. Zhao *et al.* reported that residues K133 and A176 determine MilA’s specificity for CMP over dCMP, and mutations K133R and A176S shift its specificity towards dCMP^156^. Structural alignment of MilA with the Avs trigger revealed that R139 in the Avs trigger aligns with K133 in MilA and S183 aligns with A176 (Figure 3A). These observations suggest that the Avs trigger is more likely to function as a dCMP or dUMP hydroxymethylase^28^—known to be involved in DNA hypermodification—rather than as a CMP hydroxymethylase. AlphaFold 3 also predicts a high-confidence complex between the type I Avs trigger and dUMP (ipTM = 0.97, pTM = 0.96), followed by a slightly lower confidence complexes with dCMP (ipTM = 0.94, pTM = 0.95) and CMP (ipTM = 0.92, pTM = 0.95); these scores are sufficiently similar that differences should be interpreted cautiously. While these scores are not substantially different, collectively these results suggest that the type I Avs trigger is likely to act as an ADP involved in phage DNA hyper modification.

**Figure 3.**
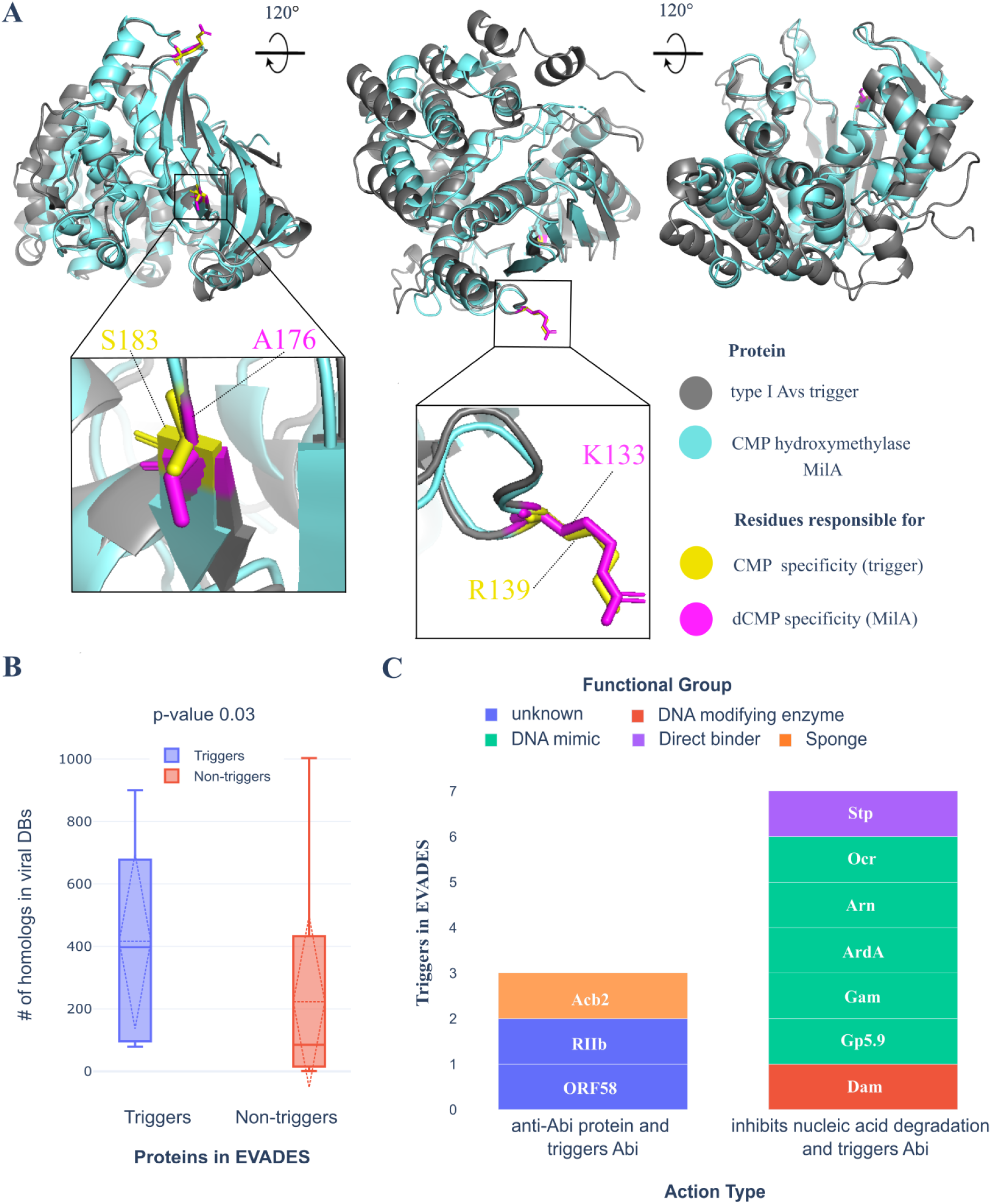
Phage ADPs trigger other defence systems. **(A)** Structural superposition of the Avs type I trigger from *Bacillus* phage SBSphiC (grey) and cytidine monophosphate hydroxymethylase MilA (cyan). Alignment was obtained with FATCAT 2.0 (p-value = 0.00^e+00^). Residues associated with CMP specificity are rendered as sticks and coloured in yellow, while those associated with dCMP specificity are in magenta and also rendered as sticks. The Avs type I trigger has substitutions likely responsible for dCMP specificity. **(B)** Boxplots showing the number of homologs of ADPs (those known to trigger bacterial defence systems [blue] vs. other ADPs [red]) in phage genomes from GenBank. **(C)** Counts of ADPs found in EVADES that trigger other defence systems, organised based on the mechanism of action (Abortive Infection, Abi) and coloured by the functional group.

An analysis in viral databases (see methods) revealed that phage ADPs reported to trigger bacterial defence systems (Supplementary Table 5) have, on average, more homologs than other ADPs (416 vs. 223, p-value 0.03, Figure 3B). These ADPs predominantly target bacterial systems that degrade phage nucleic acids and activate defences that induce abortive infection (Figure 3C). This pattern may be explained by evolutionary dynamics: bacteria are more likely to develop new defence strategies against widely prevalent phage ADPs. By coupling a system that degrades phage nucleic acids with an abortive-infection-inducing defence, bacteria can maximise their protection while minimising collateral damage to their own population due to autoimmunity, should the former fail to eliminate the phage. Further supporting this interpretation, the expression of abortive infection (Abi)-inducing defence systems, such as Gabija and Nezha, has been shown to be negatively regulated by active CRISPR systems. Inhibition of CRISPR systems leads to the overexpression of these CRISPR-supervised immune systems (CRISIS), reinforcing the model of layered immune regulation^25,157^.

Interestingly, five out of ten ADPs triggering bacterial defence systems are also DNA mimics. Given that there are 11 DNA mimic ADPs in the database (Supplementary table 6), this suggests that DNA mimics are significantly more likely to trigger bacterial defence systems (Fisher’s exact test p-value 9.53 × 10⁻⁶, Odds ratio 42), but we acknowledge the total number is small. Nevertheless, we hypothesised that DNA mimics may bind a broad range of bacterial defence proteins that target nucleic acid^99^, placing evolutionary pressure on bacteria to develop counter-counter adaptations—defence systems capable of recognising them. To investigate this, we co-folded DNA mimics with bacterial defence systems and predicted 20 high-confidence interactions (Supplementary Table 7), and 10 out of 20 of these interactions were with bacterial defence proteins binding DNA and 9 out of 20 were with the effector proteins. Among the 9 effectors, 4 have uncharacterised mechanisms of action, while 3 target DNA: BrxC, which binds host DNA as part of the BrxBCXZ complex in the BREX defence system^14^; HamA, the nuclease effector of the Hachiman defence that degrades both host and phage DNA^158^; and KwaB, which binds phage DNA to block replication in the Kiwa defence^159^. Unexpectedly, the remaining 2 effectors were cyclases from type I and type IV CBASS defence systems.

### Eukaryotic viruses encode several homologs of phage ADPs

In order to assess conservation of defence evasion strategies between bacterial and eukaryotic viruses, we looked for structural homologs of phage ADPs in a database of ∼68,000 proteins encoded by eukaryotic viruses, including ∼43,000 proteins originating from dsDNA viruses^160^. We identified structural homologs for 27 ADPs, predominantly anti-RM proteins (n=10), followed by Acrs (n=5), anti-Avs (n=2) proteins, and anti-PrrC proteins (n=2). Interestingly, several of these ADPs evade defence systems that are evolutionarily conserved between bacteria and eukaryotes^32,161,162,163^. For instance, the phage encoded ORF55 and ORF83 proteins evade the bacterial type III Avs (antiviral STAND) defence system^17^, which shares homology with the STAND superfamily of pattern-recognition receptors involved in eukaryotic innate immunity^32^. Similarly, phage encoded nucleotidyltransferases (NTases) inhibit the STING pathway, a well-known immune signalling mechanism found in both bacteria and eukaryotes^53^. We next conducted a detailed analysis of these structural similarities to evaluate whether the underlying mechanisms of action are also shared between phage and eukaryotic viruses.

### Anti-Avs ORF55 is a DNA ligase shared by bacterial and eukaryotic viruses

ORF55 is an anti-Avs type III protein of phage *DruSM1*^17^. Bacterial Avs systems detect phage-encoded proteins such as terminases and portal proteins, triggering homo-oligomerisation and subsequent activation of the N-terminal effectors domains. Among these, DNA nucleases are the most prevalent effectors, inducing non-specific DNA cleavage upon activation. Interestingly, it has been demonstrated that bacterial Avs can also recognise portal proteins and terminases from eukaryotic viruses, including those from human herpesvirus 8^32^.

ORF55 has been annotated as a DNA ligase^17^. Based on this annotation, we hypothesised that it may counteract the non-specific dsDNA endonuclease^32^ activity of the type III Avs effector proteins by ligating DNA fragments cleaved during Avs activation. To investigate this possibility, we searched for structural homologs of ORF55 in PDB. The closest match was the *Chlorella* virus DNA ligase (PDB: 2Q2U, p-value = 0.00^e+00^)^164^, a well-characterised enzyme involved in DNA damage repair^165^ (Figure 4A). This DNA ligase represents the minimal functional unit of the ATP-dependent DNA ligases and exhibits intrinsic nick-sensing activity, binding with high affinity to duplex DNA containing a single 3′-OH–5′-PO4 nick^166^. Structural alignment revealed that ORF55 retains 11 out of 13 active site residues conserved in the *Chlorella* virus DNA ligase (Figure 4B)^164^. We used AlphaFold 3 to model the structure of ORF55 in the complex with the DNA fragment co-crystallised with the *Chlorella* virus DNA ligase, which produced a high-confidence complex (ipTM 0.88, pTM = 0.91) with a similar DNA binding interface (Figure 4A). Given that the *Chlorella* virus ligase efficiently ligates nicked DNA, performs poorly on one-nucleotide gaps, and fails entirely on two-nucleotide gaps^166^, we modelled three complexes: ORF55 with ATP and (i) nicked DNA, (ii) one-nucleotide gapped DNA, and (iii) two-nucleotide gapped DNA. AlphaFold 3 predicted with higher confidence the nicked (ipTM = 0.95, pTM = 0.95) and one-nucleotide gapped complexes (ipTM = 0.95, pTM = 0.95) than the two-nucleotide gapped complex (ipTM = 0.87, pTM = 0.91). These results suggest that ORF55 from the DruSM1 phage shares a similar substrate preferences to the *Chlorella* virus DNA ligase, favouring nicked and one-nucleotide gapped DNA.

**Figure 4.**
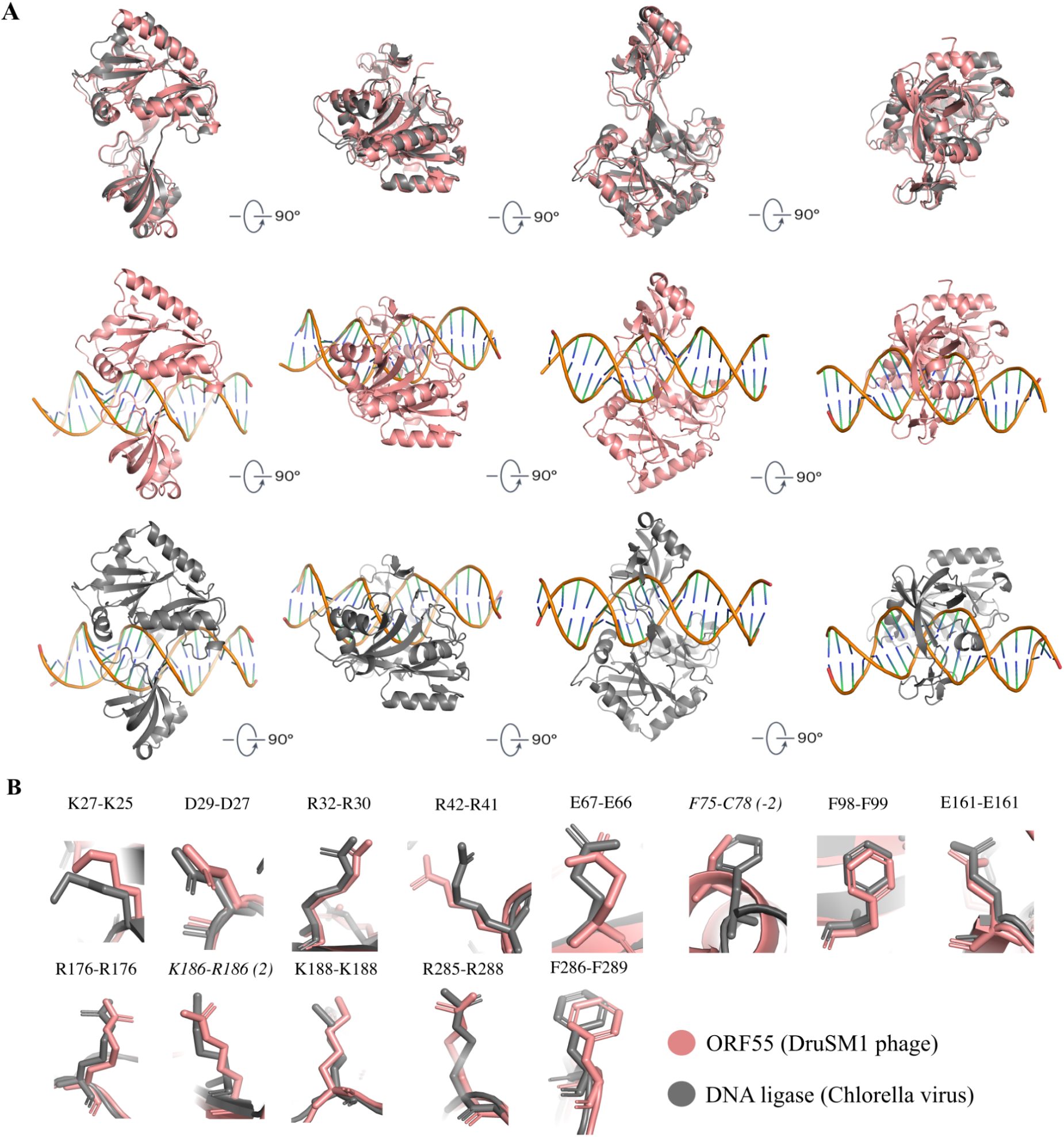
ORF55 shares high similarity with the DNA ligase from Chlorella virus. **(A)** From left to right, each view represents a 90° rotation of the previous view around the x-axis. Top: superposition of the *Chlorella* virus DNA ligase (salmon, ribbon representation, PDB: 2Q2U) and ORF55 (grey, ribbon representation) generated using FATCAT 2.0 (p-value 0.00^e+00^). Middle: the DNA ligase from *Chlorella* virus in complex with DNA (PDB: 2Q2U). Button: a high-confidence complex between ORF55 and the DNA fragment from the complex with the DNA ligase from *Chlorella* virus. **(B)** Thirteen residues contributing to the active site of DNA ligase (grey) are rendered as sticks aligned with the closest residues (rendered in the same way) from ORF55 (salmon).

We identified homologs of ORF55 in 66 different species of eukaryotic dsDNA viruses infecting a wide range of hosts, including amoebae, algae, insects, invertebrates, birds, and mammals, most notably humans. The ORF55 homologs that have been functionally characterised and are involved in DNA repair, for example, the eukaryotic African swine fever virus (ASFV) encodes a homologous DNA ligase. ASFV replicates and assembles virions in the cytoplasm of infected cells, primarily macrophages, which are highly enriched in reactive oxygen species (ROS) and subject to extensive DNA damage. The ASFV DNA ligase has been reported as a crucial component of its DNA repair system^167^. Although viral DNA ligases are generally non-essential for eukaryotic virus DNA replication, since these viruses can utilise cellular DNA ligases for replication, they are essential for conferring resistance to DNA-damaging agents^168^. Normally, the cytosol of eukaryotic cells does not contain dsDNA, and several nucleases act to degrade any cytosolic DNA that arise^169–171^. Therefore, maintaining genome integrity is extremely important for DNA viruses, as DNA damage can trigger antiviral responses such as interferon (IFN)^172^ or inflammasome activation^173^, ultimately limiting viral propagation.

### Anti-Avs protein ORF83 is likely to bind AMP and is encoded by viruses infecting multiple amoeba species

ORF83 is an anti-type III Avs protein, originally annotated as “Smf; Predicted Rossmann fold nucleotide-binding protein DprA/Smf involved in DNA uptake”^17^. We predicted a high-quality structural model of ORF83 using AlphaFold 3 (pLDDT 95, pTM = 0.92) and searched for structural homologs in PDB^174^. The top-scoring matches were *Cg*1261 (PDB ID: 5WQ3, E-value = 4.39e^-7^), a type II cytokine-activating protein from the Lonely Guy (LOG) family, along with TT1465, a hypothetical lysine decarboxylases (PDB ID: 1WEK, E-value = 7.95e^-7^), and DprA (PDB ID: 3UQZ, E-value = 3.49e^-5^). Although TT1465 was initially predicted to be a lysine decarboxylase, experimental studies revealed that it lacks decarboxylase activity and instead exhibits phosphoribohydrolase activity^175^. Both *Cg*1261 and TT1465 were shown to hydrolyse AMP into adenine and ribose 5-phosphate^175^. However, among these, only *Cg*1261 has been experimentally confirmed to possess cytokine-activating activity and the ability the ability to convert N6-(isopentenyl)-adenosine 5′-monophosphate (iPRMP) into isopentenyladenine via dephosphoribosylation.

To investigate whether ORF83 might have similar phosphoribohydrolase activity, we modelled its interaction with AMP using AlphaFold 3, yielding a high-confidence complex (ipTM = 0.9, pTM = 0.88; Figure 5A). Notably, the AMP-binding region of ORF83 aligns with the corresponding AMP-interacting region of *Cg*1261 (Figure 5A). Structural alignment between the active site of *Cg*1261 and ORF83 revealed that they share two invariant residues (R115/R89 and T175/T112), along with a conservative (E178/D114) and a neutral substitution (K116/N90). Given the high structural similarity between ORF83 and *Cg*1261, the confidently predicted AMP interaction, and the partial conservation of active site residues, we propose that ORF83 likely possesses phosphoribohydrolase activity towards AMP. However, the residue composition of ORF83 diverges from that of *Cg*1261 at the prenyl-binding site, thus providing no structural evidence that ORF83 binds iPRMP or functions as a cytokine-activating protein.

**Figure 5.**
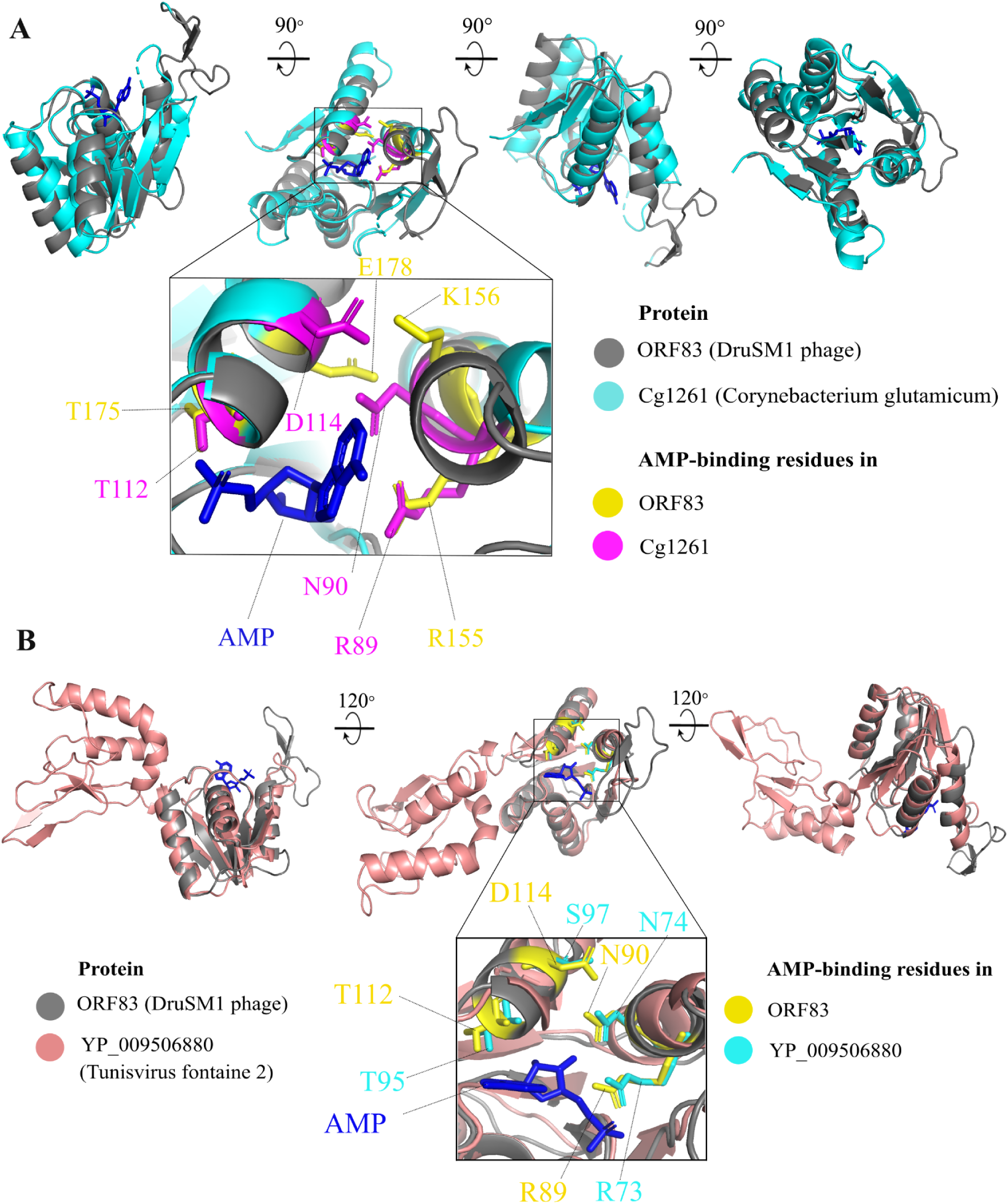
ORF83 is a homolog of Cytokine-Activating Protein Cg1291, binds AMP, and shares similarity with proteins in eukaryotic dsDNA viruses. **(A)** Four different views of the superimposition of the AMP (blue, stick representation)-ORF83 (grey, ribbon representation) complex (AlphaFold 3 prediction) with the type II cytokine-activating protein *Cg*1261 (cyan, ribbon representations) of the LOG family, generated using FATCAT 2.0 (p-value = 4.39^e-7^), each orientation is related by a 90° rotation around the x-axis. The second view provides the best orientation of the AMP-binding site, which is expanded in the bottom insert. Residues in *Cg*1261, characterised as part of the AMP-binding active site, are coloured magenta, and the closest residues in ORF83 are coloured yellow (corresponding to AMP-binding residues predicted by AlphaFold 3). **(B)** Three different views of the superimposition of the complex between AMP (blue, stick representation) and a protein from *Tunisvirus fontaine 2* (salmon, ribbon representation, NCBI GenPept accession: YP_009506880) with the phage protein ORF83 (grey, ribbon representation), generated using FATCAT 2.0 (p-value = 2.99^e-13^) each related by a 120°.

We examined the weaker structural similarity between ORF83 and DprA, but ORF83 lacks the equivalent key residues involved in DprA self-dimerisation and RecA binding^176^. AlphaFold predicts a low-likelihood interaction with RecA (ipTM = 0.18, pTM = 0.61). Instead, ORF83 likely binds AMP and functions as a phosphoribohydrolase, though experimental validation is needed to confirm this and clarify its role in evading type III Avs defence.

We found 14 homologs of ORF83 in eukaryotic viruses that all infect various amoeba species. Interestingly, many homologs are longer than ORF83 and contain the domain DUF4326 at the C-terminus. For instance, YP_009506880 from *Tunisvirus fontaine 2* shares extremely high structural similarity to ORF83 (p-value = 2.99^e-13^; Figure 5B). We predicted its complex with AMP using AlphaFold 3 (ipTM = 0.85, pTM = 0.77). The alignment between ORF83 and YP_009506880 showed that they share the same putative AMP-binding active site (R89-R73, N90-N74, and T112-T95 in ORF83 and YP_009506880, respectively), differing only by one neutral substitution (D114-S97). Overall, these results demonstrate functional homology between ORF83, the anti-Avs protein from phages, and YP_009506880, a protein from eukaryotic viruses.

### Phage nucleotidyltransferase homologs in eukaryotic viruses

The cyclic GMP–AMP synthase (cGAS)-STING pathway is a conserved immune system in bacteria and eukaryotes. It is characterised by sensory cGAS proteins that synthesise cyclic nucleotides in response to viral infections. These nucleotides bind the STING domain-containing effector proteins, which initiate downstream immune responses. In bacteria, these effectors typically cleave NAD+, whereas in eukaryotes they activate the expression of antiviral genes.

An NTase enzyme from *Bacillus* phage Bcp1 (NCBI GenPept accession: YP_009031408) synthesises a competing cyclic nucleotide, 3′,3′-cyclic GMP-AMP (3′,3′-cGAMP), which inhibits the cGAS-STING response^53^. We identified 12 homologs of this phage NTase in a range of eukaryotic viruses that infect insects, suggesting a potential role in suppressing the eukaryotic cGAS-STING pathway. For instance, YP_009091875 from *Erinnyis ello* granulovirus shows high structural similarity to the phage NTase (Figure 6A; FATCAT 2.0 p-value 3.10^e-10^). Importantly, Ho *et al.* identified two catalytic motifs for phage NTases: GS-x-AY[GAN]T-x4-SDxD and NP-x-h2[DE]. The second motif (NP-x-h2[DE]) is unique to phage NTases within the broader NTase superfamily. Notably, the *Erinnyis ello* granulovirus protein exhibits comparable similarity to the consensus active site motifs as the Bcp1 NTase (Figure 6B), matching 10 out of 14 conserved positions, whereas the phage NTase with experimentally verified inhibitory activity against cGAS-STING response matches 11 out of 14.

**Figure 6.**
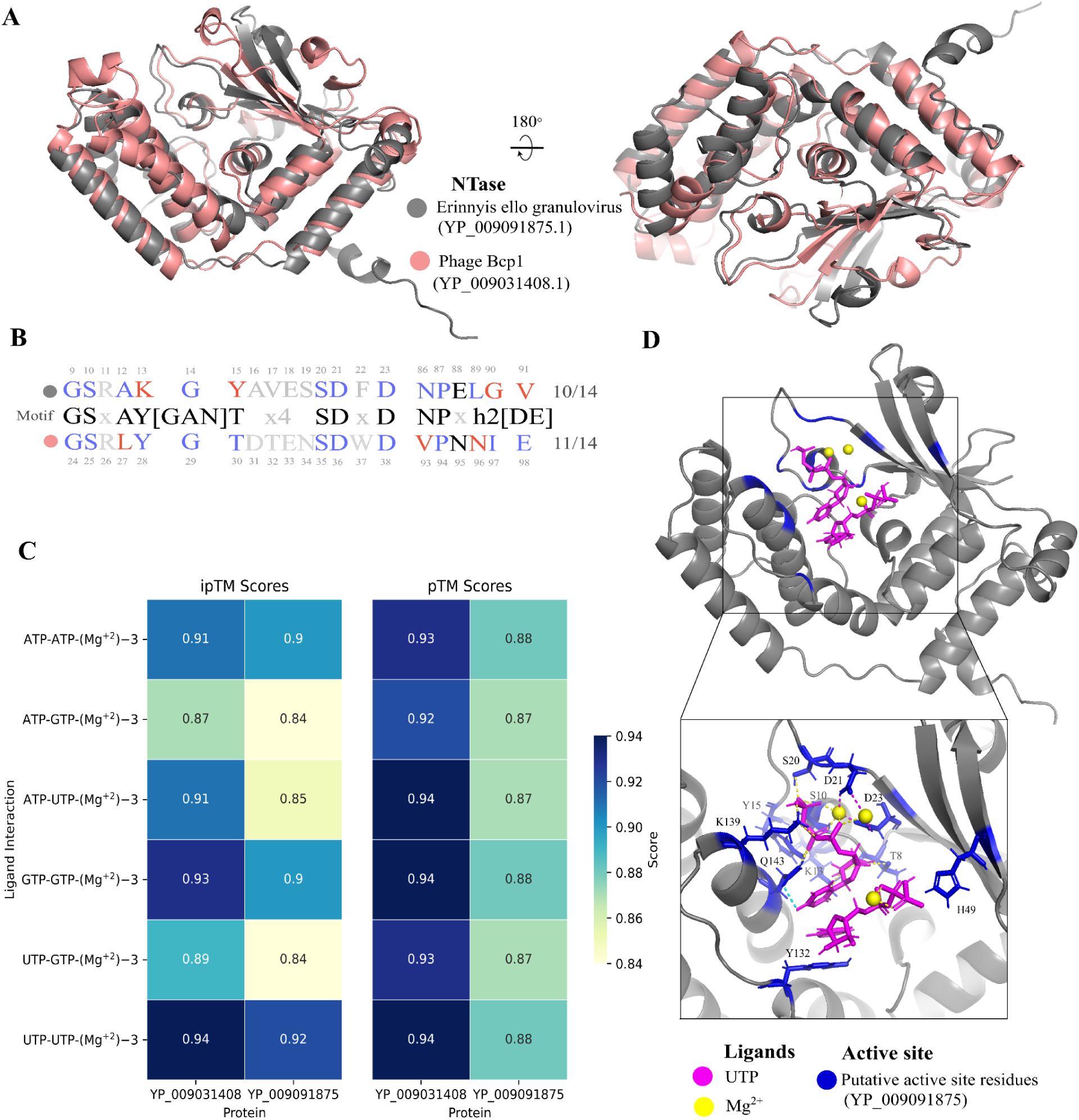
Structural and Functional Analysis of NTase from Bacillus Phage Bcp1 and Erinnyis ello Granulovirus. **(A)** Left, structural alignment of the cyclic nucleotide-synthesising NTase from *Bacillus* phage Bcp1 (salmon, NCBI GenPept accession: YP_009031408) and its homolog from *Erinnyis ello* granulovirus (grey, NCBI GenPept accession: YP_009091875), performed using FATCAT 2.0 (p-value: 2.74^e-10^). Right, the view of the structural alignment is rotation 180° around the X-axis. **(B)** Alignment of catalytic site motifs of phage NTases (middle row) to the experimentally validated NTase from *Bacillus* phage Bcp1 (bottom row) and its homolog from the dsDNA eukaryotic virus (YP_009091875 from *Erinnyis ello* granulovirus, top row). Matches are in blue, mismatches in red, and residues matching variable motif positions in the sequence motif are in grey. **(C)** Heatmap of confidence scores (ipTM and pTM) for complexes between phage or viral NTases with pairs of NTPs and three ions of Mg^2+^ produced by AlphaFold 3. **(D)** Predicted complex of *Erinnyis ello* granulovirus NTase (grey) with two molecules of UTP (magenta) and three ions of Mg2+ (yellow), the active site residues in the NTase are in blue.

Nucleoside triphosphates (NTPs) serve as ligands in cyclic nucleotide synthesis, with metal ions such as Mg2+ typically acting as catalytic cofactors^177^. To investigate functional mechanisms of these NTases further, we modelled interactions between NTPs used in cyclic nucleotide synthesis (ATP, GTP and UTP) with both the phage NTase and its homolog in *Erinnyis ello* granulovirus using AlphaFold 3. All predicted complexes yielded ipTM scores above 0.8 (Figure 6C, Supplementary Table 8), indicating high-confidence predictions, and the predicted ipTM scores for complexes of phage and viral proteins show strong correlation (Pearson correlation = 0.865, p-value 0.0261), likely indicating similar substrate preferences. The highest-scoring complex produced for both proteins includes two molecules of UTP and three ions of Mg^2+^. We show that the highest-confidence complex predicted for protein from *Erinnyis ello* granulovirus uses a similar configuration of the active site as *E. cloacae* CD-NTase CdnD (*Ec*CdnD), in which two Mg²⁺ ions coordinate the donor NTP and one Mg²⁺ ion coordinates the acceptor NTP (Figure 6D)^177^. Moreover, we visualised the residues forming bounds with NTPs and ions and showed that conserved motif SDxD is particularly important, similar to the E-clade of CD-NTases^177^ (Figure 6D).

Given the high structural similarity between the phage NTase and putative NTases from the dsDNA eukaryotic viruses, their conserved active site residues, and high-confidence predictions of protein-NTPs-Mg2+ complex formation, we postulate that the viral proteins also synthesise cyclic nucleotides. This activity may enable eukaryotic dsDNA viruses to evade the cGAS-STING response.

Additionally we examined matches between proteins in EVADES and IMG/VR database and identified homologs of Tad1 in three high-confidence viral genome fragments assigned to the order *Durnavirales*. Members of this order are double-stranded RNA (dsRNA) viruses known to infect eukaryotic hosts and these fragments encode RNA-dependent RNA polymerase (RdRp) which serves as a hallmark of RNA viruses. Given that Tad1 can sequester universal cyclic nucleotides utilised by both bacterial CBASS and eukaryotic cGAS immune pathways, such as 2′,3′-cGAMP^178^, we speculated that these viral homologues may contribute to evasion of eukaryotic cGAS-mediated immunity. To investigate this hypothesis, we modelled the complex between the *Durnavirales*Tad1 and 2′,3′-cGAMP and it yelled a high-confidence complex (ipTM = 0.90, pTM = 0.91). Structural comparison of the predicted DurnaviralesTad1–2′,3′-cGAMP complex with the experimentally determined Tad1–2′,3′-cGAMP structure (PDB ID: 8KBG) revealed an almost identical overall architecture (Supplementary Figure 4), indicating a conserved mechanism of cyclic dinucleotide recognition. However, because these homologs derive from metagenomic fragments rather than complete viral genomes, definitive evidence that *Durnavirales* species employ Tad1-like proteins to subvert host immunity remains to be established.

### Putative ADPs acting via direct binding from phage diversity hotspots

Diversity hotspots are genomic regions marked by elevated sequence variability, typically flanked by conserved regions or genes, and commonly found in phage and plasmid genomes. These regions emerge through mechanisms such as mobile genetic element insertions, recombination, and hypermutation, and are often enriched in accessory genes related to defence, anti-defence, virulence, and antimicrobial resistance^179^. Notably, similar to bacterial defence systems, phage anti-defence genes frequently co-localise within these hotspots^7,46,53^. We hypothesised that many uncharacterised proteins encoded near anti-defence genes in phage diversity hotspots may contribute to immune evasion. It has been demonstrated^37^ that AlphaFold 2 Multimer^180^ can predict phage proteins that directly bind and inhibit bacterial defence systems, validating anti-defence activity in 6 out of 21 predicted interactors. Building on this approach, we developed a similar pipeline based on AlphaFold 3 (Supplementary Figure 5), which outperforms AlphaFold 2 in predicting protein-protein interactions^38^, to identify phage proteins that interact with bacterial defence systems—focusing specifically on uncharacterised proteins prevalent in diversity hotspots. Using this pipeline, we conducted a large-scale screening of proteins with unknown function from phage diversity islands and identified 46 predicted to bind bacterial defence proteins with high confidence. These predicted ADPs target proteins from 27 distinct families of bacterial defence systems, with ISG15-like systems being the most frequently targeted. As interactions with effector proteins are more likely to result in defence inhibition, we prioritised candidates predicted to bind such effectors, yielding 29 proteins targeting 15 different defence systems (Supplementary Table 9). Yirmiya *et al.* reported that AlphaFold predictions are more reliable when remote homologs are also predicted to bind the same target protein. Accordingly, we co-folded remote homologs of phage proteins with their corresponding defense proteins; two of these pairs passed the threshold confidence score of 0.8. We then incorporated these results to refine the prioritisation of candidate ADPs for future experimental validation.

## DISCUSSION

We have curated a comprehensive database of phage ADPs, called EVADES. This database encompasses a larger diversity of ADPs than other resources, such as Anti-DefenseFinder^147^ or dbAPIS^148^. More importantly, it provides insights into the MoA of the APDs, as well as catalogues both experimental and predicted structures, information about protein family membership, and the potential to trigger bacterial defence systems. We also categorised phage ADPs based on their MoA. While most ADPs remain uncharacterised, the most prevalent functional characterised category is direct binders–phage proteins that directly interact with bacterial defence proteins to inhibit their activity. We believe this functional information can facilitate biological insights. For instance, knowing that the majority of known ADPs are direct binders, we hypothesised that some uncharacterised ADPs may inhibit bacterial defences through binding. To test this hypothesis, we modelled the interactions between uncharacterised ADPs and the defence systems they inhibit using AlphaFold 3, and predicted interactions for 20 ADPs. Where possible, we cross-referenced the predicted complexes and experimental data and found that our predictions align with the experimental results. One exciting example is AcrIB4, a protein that was reported as a Cas8b homolog (sharing 38% identity with the last 90 residues), but was not replacing Cas8b in the Cascade complex^58^. We predicted that AcrIB4 binds to Cas8b, specifically interacting with the residues in Cas8b that bind to the Cas11 protein, which stabilises the Cascade complex. Cas11 is translated from an internal ribosome-binding site within the 3′ end of the *cas8b* gene, suggesting that AcrIB4 is homologous to Cas11. We propose that AcrIB4 is likely to replace Cas11, destabilising the Cascade complex. To the best of our knowledge, this is the first example of an Acr acting through complex subunit poisoning by replacing Cas11.

Many ADPs, while inhibiting one defence system, may actually trigger another. These ADPs are widespread and inhibit ubiquitous defence systems including those that cleave phage genetic material. At the same time, those may trigger a secondary line of defences that induce abortive infection. We speculate that this could be a common phenomenon in phage–host interactions, as bacteria evolve counter-counter-defence systems in response to phage counter-defence strategies. Furthermore, we hypothesise that bacterial defence systems operate in layers, where systems that cleave foreign nucleic acids and cause minimal collateral damage serve as the first line of defence. In contrast, systems that trigger abortive infection may sense the inhibition of these first-line defences, providing maximum protection while minimising self-damage. Such interactions between defence systems may also explain the distribution of these systems in bacterial populations, where certain pairs are frequently encoded together due to their evolutionary advantages^181,182^. We showed that ADPs that trigger bacterial defence systems are enriched in those ADPs that act as DNA mimics. We predicted broad-spectrum direct interactions between DNA mimics and bacterial defence proteins, unsurprisingly, particularly those that bind DNA. This layering of defence systems could be explained by the evolutionary pressure exerted by DNA mimics on bacteria, as they can inhibit numerous DNA-binding defence systems, thereby driving the evolution of bacterial counter-counter adaptations. Finally, we found that two proteins reported previously as triggers of bacterial defence systems are identical to ADPs in our database.

We identified numerous homologues of phage ADPs in eukaryotic viruses, suggesting that immune evasion strategies can be conserved across bacterial and eukaryotic viruses. For example, eukaryotic dsDNA viruses encode structural and functional homologues of phage nucleotidyltransferases, which in phages inhibit the bacterial STING effector by producing competing cyclic nucleotides. Because the cGAS–STING pathway is also conserved in eukaryotes, these competing oligonucleotides could similarly suppress the eukaryotic cGAS–STING response—a key antiviral defense—thereby facilitating viral infection. This conservation also provided mechanistic insights into the activity of the phage anti-Avs protein ORF55, informed by its similarity to the minimal nick-sensing DNA ligase found in *Chlorella* virus. Together, these findings highlight how studies of phages can illuminate the immune evasion strategies of eukaryotic viruses, underscoring the potential for reciprocal discovery between bacterial and eukaryotic viral systems.

Finally, we conducted a large-scale computational screen to identify new ADPs among uncharacterised phage proteins, using a previously established approach and focusing on protein encoded in phage diversity islands with anti-defence genes. Thereby, we predicted 29 new candidate ADPs targeting effector proteins of bacterial defence systems. Altogether, EVADES not only consolidates known ADPs into a single, richly annotated resource, but also facilitates elucidation of their mechanisms of action, exploration of their evolutionary dynamics, and discovery of new ADPs.

## Limitations of the study

Our resource paper provides a comprehensive collection of phage ADPs and proposes several hypotheses which are based on in-depth computational analysis, but are yet to be supported by experimental validation. To enable the users of EVADES to understand those functions that are predicted versus experimentally validated, MoA are labelled with evidence code ontologies. In this paper and the resource, we have focused on high-confidence predictions, reducing the chances of false positives. The analysis in the section devoted to ADPs acting as triggers is based on a small number of entries from the literature, and while the results we present are statistically significant, they may change as more data become available. Although we demonstrated that ORF83 likely binds AMP and has a conserved structure between bacterial and eukaryotic viruses, we have yet to establish the detailed MoA to substantiate a claim that the homolog in eukaryotic viruses also plays a role in immune evasion. Only two out of twenty-nine predicted direct binders maintained high confidence scores (>0.8) in the analysis conducted for remote homologs, suggesting that some of the predictions could be false positives.

---

**Supplementary Figure 1.**
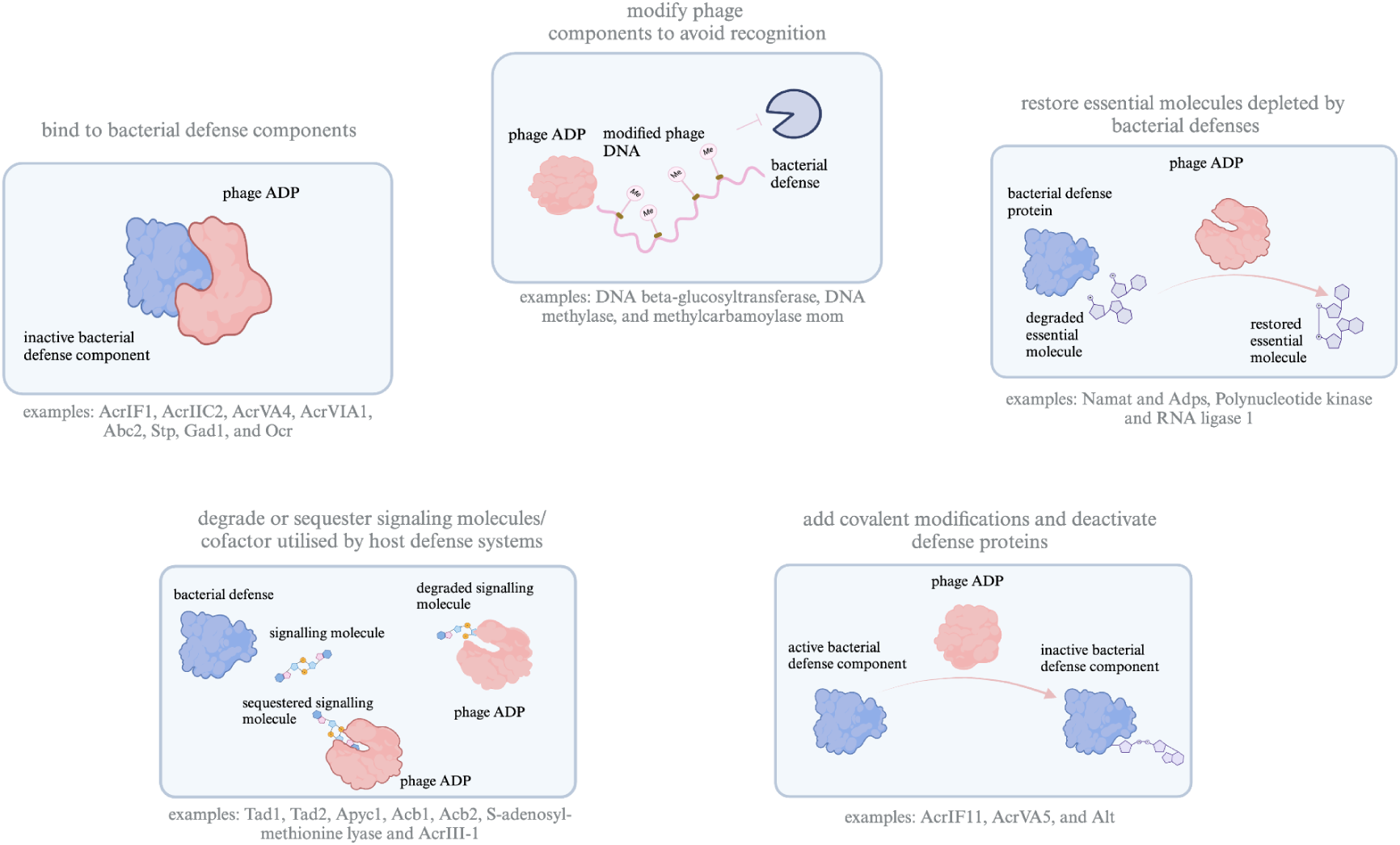
Classification of phage ADPs based on mechanism of action. ADPs can be categorised based on their mechanisms of action against bacterial defence systems: (a) ADPs that directly bind and inactivate bacterial defence components (e.g., AcrIF1, AcrIIC2, AcrVA4); (b) ADPs that degrade or sequester signalling molecules or cofactors required by host defences (e.g., Tad1, Acb1, AcrIII-1); (c) ADPs that modify phage components such as DNA to evade recognition (e.g., DNA beta-glucosyltransferase, methylases); (d) ADPs that restore essential molecules depleted by bacterial defences (e.g., Namat, Adps, RNA ligase 1); and (e) ADPs that add covalent modifications to deactivate defence proteins (e.g., AcrIF11, AcrVA5, Alt).

**Supplementary Figure 2.**
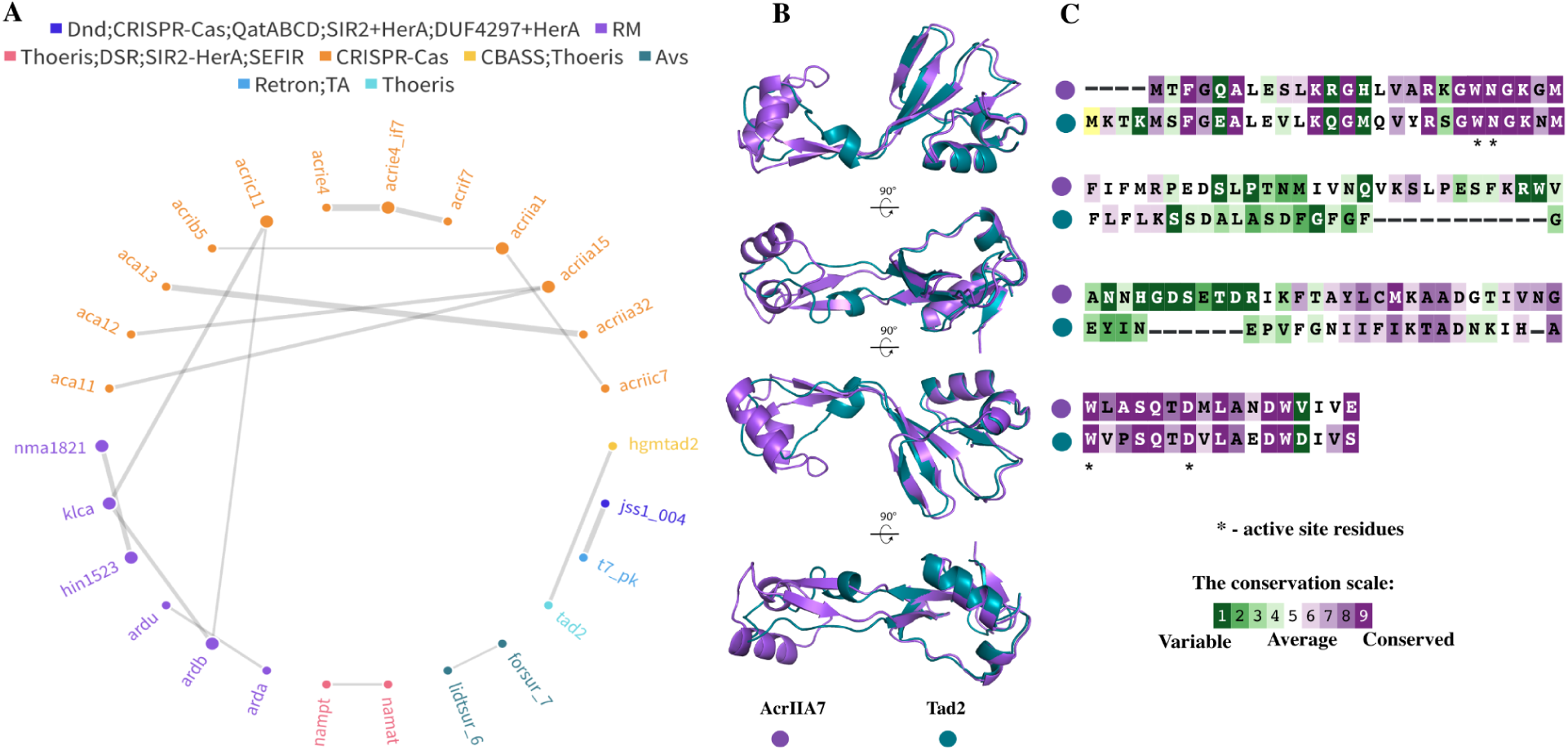
Similarity between AcrIIA7 and Tad2. **(A)** ADP sequence similarity network, ADP node size is proportional to number of proteins an ADP shares similarity to and nodes are coloured based on the defence system(s) they inhibit, edge width is proportional to shared sequence identity. **(B)** Structural alignment between AcrIIA7’s structural model (purple) produced by AlphaFold 3 and Tad2 (teal blue, PDB ID: 8SMG) obtained with FATCAT 2.0 (p-value: 2.55^e-8^). From top to bottom, each view represents a 90° rotation of the previous view around the Y-axis. **(C)** Sequence alignment between AcrIIA7 and Tad (based on structural alignment produced by FATCAT 2.0), residues are coloured based on conservation scores produced by Consurf. Active site residues denoted by asterisk (*).

**Supplementary Figure 3.**
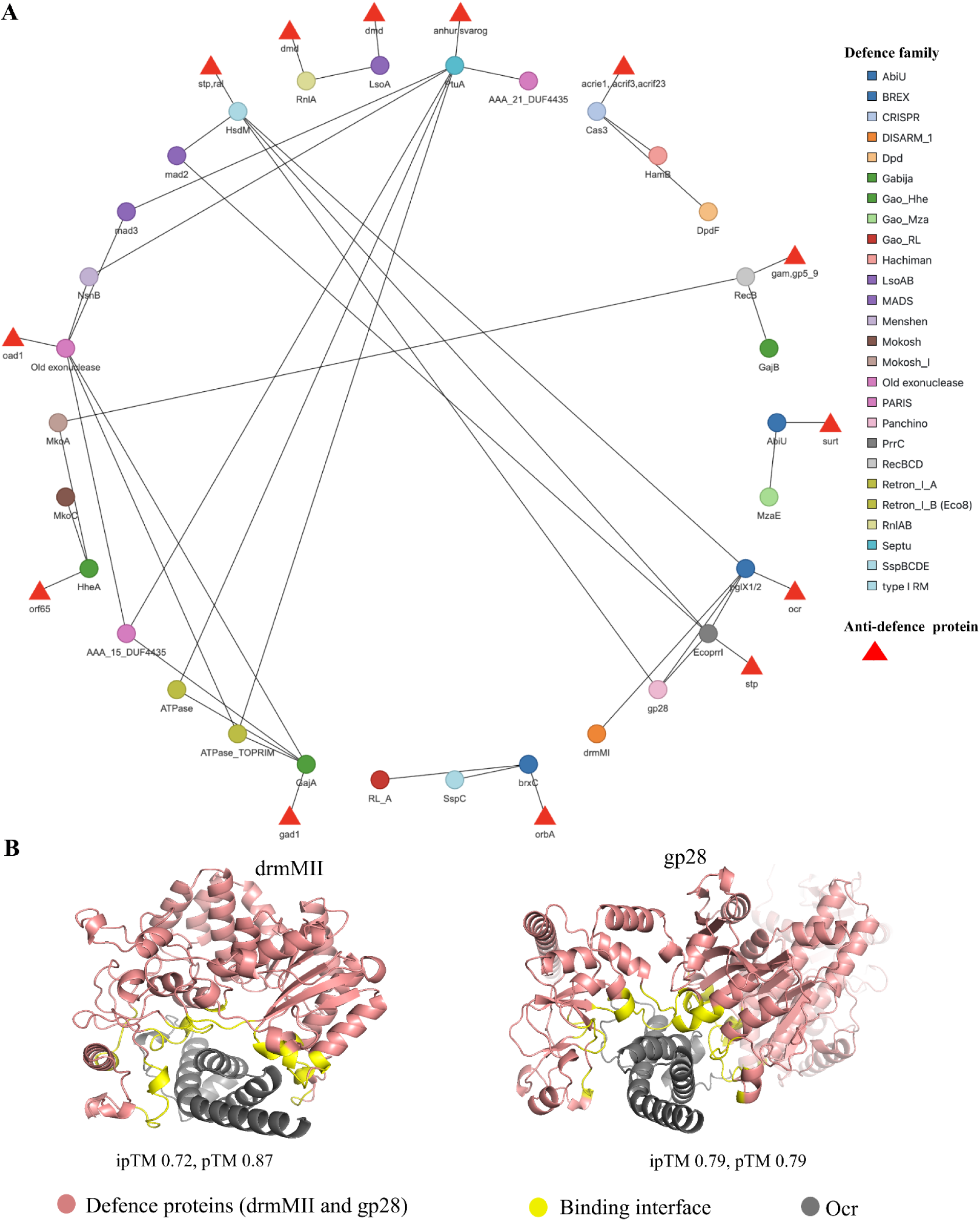
Defence protein sharing can mediate multi-defence inhibitory activity of ADPs. **(A)** Defence protein (circle nodes, coloured according to the defence system family) similarity network, where edges denote protein pairs that have similar structural folds (FoldSeek e-value = 1.0^e-8^). ADPs inhibiting the defence system directly or inhibiting single-protein defence systems are denoted as triangular nodes and are connected to the defence protein they target by an edge. **(B)** Complexes Ocr(grey)-drmMII(salmon) and Ocr(grey)-gp28(salmon) predicted by AlphaFold 3. Binding interfaces in drmMII/gp28 are shown in yellow.

**Supplementary Figure 4.**
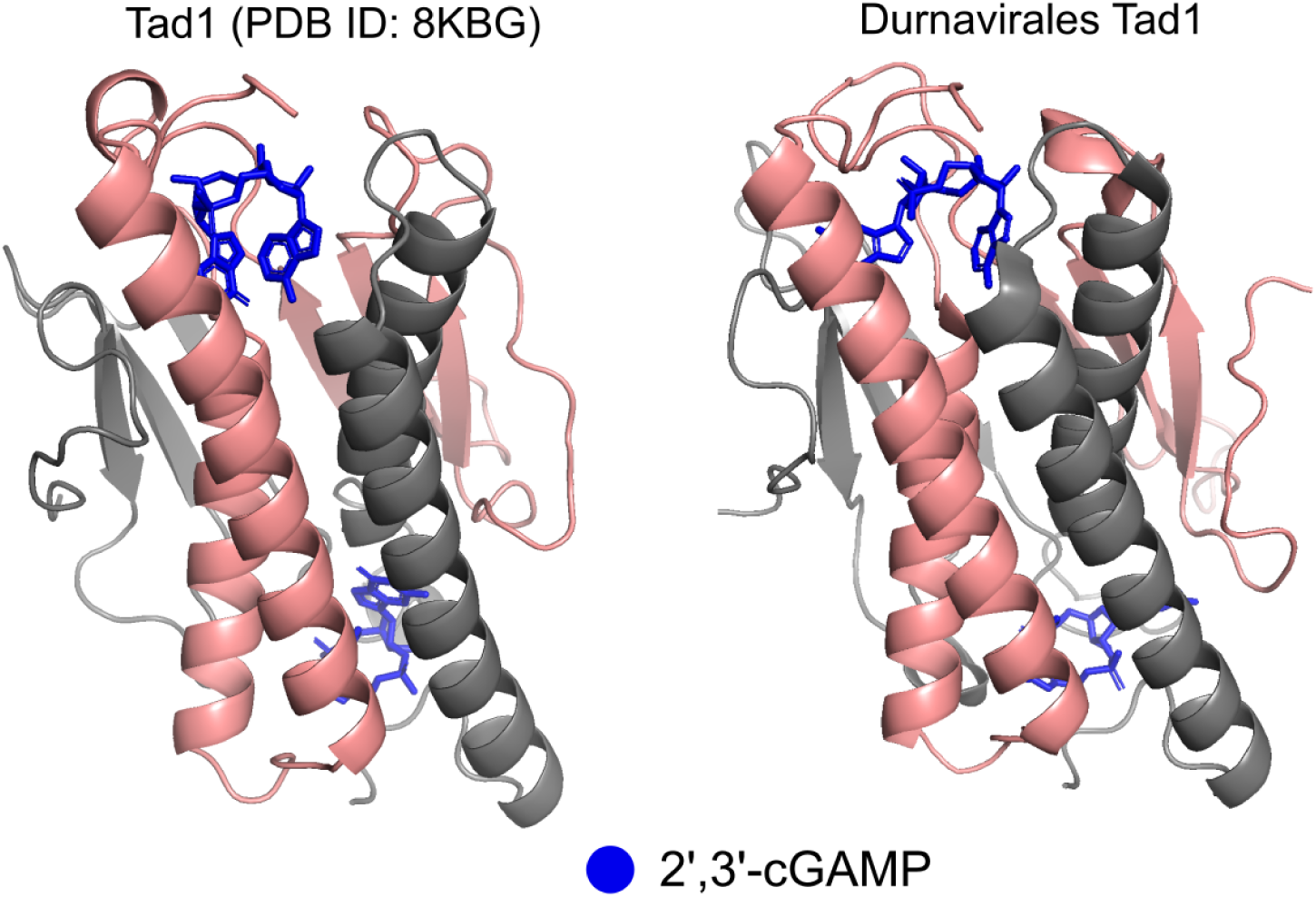
Genome fragments assigned to order Durnavirales (eukaryotic dsRNA viruses) encode homologs of Tad1. Experimentally determined structure of the complex between a Tad1 dimer (salmon pink and grey) and two molecules of 2’,3’-cGAMP (blue) is shown on the left. The predicted high-confidence complex (ipTM = 0.90, pTM = 0.91) between a dimer of the Tad1 homolog encoded in Durnavirales (IMG/VR UViG: IMGVR_UViG_3300019224_000070, salmon pink and grey) and two molecules of 2’,3’-cGAMP is shown on the right.

**Supplementary Figure 5.**
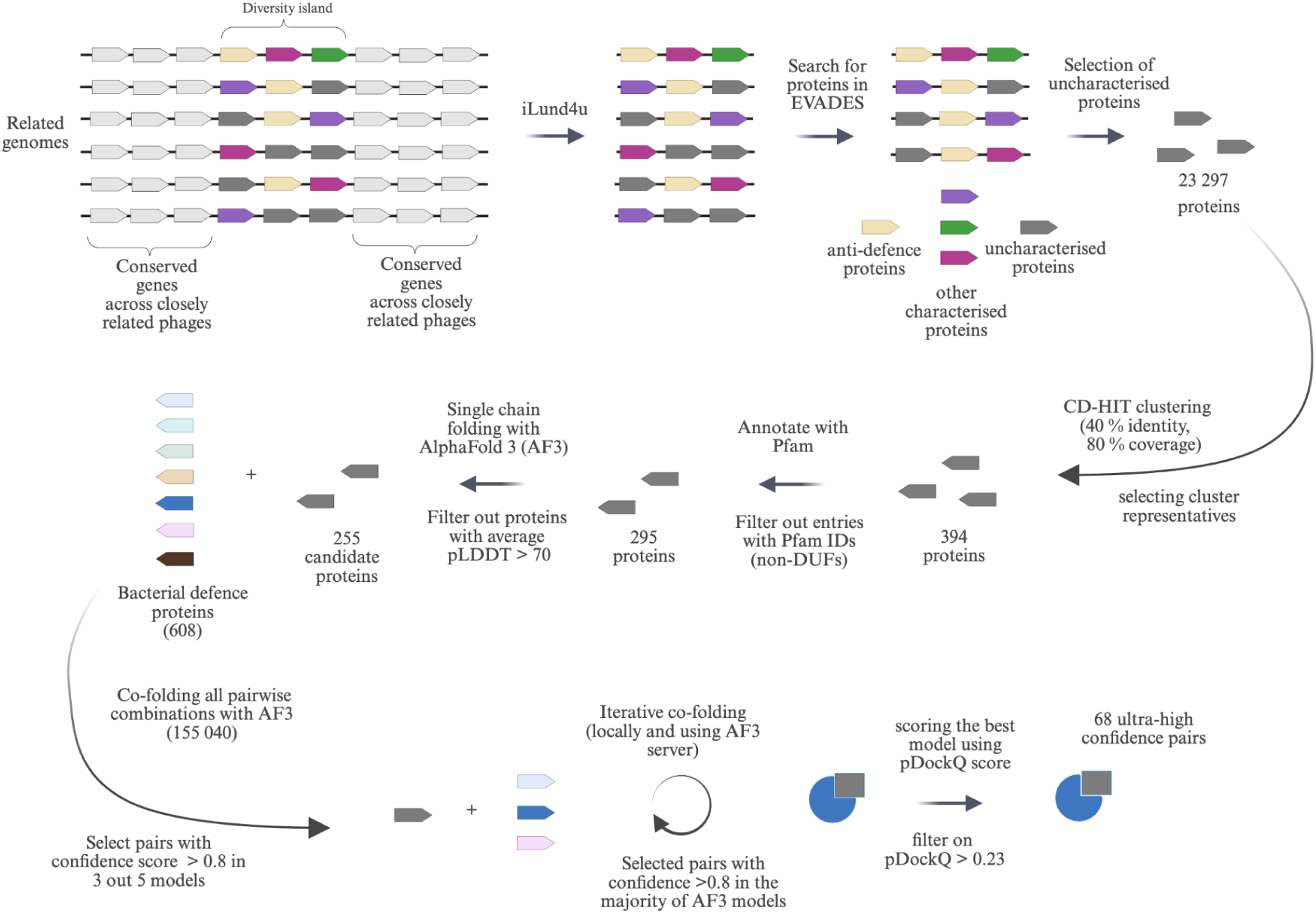
Computational pipeline used for predicting proteins binding bacterial defence systems. The co-folding pipeline selects uncharacterised protein clusters from phage diversity hotspots, filters out those with Pfam annotations or low-confidence models, and applies iterative co-folding to identify interactions with bacterial defences. Phage–bacterial protein pairs were modelled with AlphaFold3, and only those consistently reaching a high confidence score were retained. Finally, interactions were validated with pDockQ, keeping only pairs with scores above 0.23.

## RESOURCE AVAILABILITY

### Lead contact

Further information and requests for resources should be directed to and will be fulfilled by the lead contact, Robert D. Finn.

### Materials availability

Materials generated in this study have been deposited to: https://ftp.ebi.ac.uk/pub/databases/metagenomics/research-team/projects/evades/

### Data and code availability

The database is publicly available at: https://www.ebi.ac.uk/finn-srv/evades

## ACKNOWLEDGMENTS

K.M., F.C, and R.D.F. are funded by the European Molecular Biology Laboratory (EMBL). E.K. is an ARISE Fellow and received funding from the European Union’s Horizon 2020 research and innovation programme under the Marie Skłodowska-Curie grant agreement No 945405. J.W. was a recipient of Marie Skłodowska-Curie Postdoctoral Fellowship, EMBL Interdisciplinary Postdoctoral (EIPOD) Fellowship and EMBO Non-Stipendiary Fellowship (EMBO ALTF 400-2022), and a Junior Research Fellow at Wolfson College, the University of Cambridge, UK.

We would like to thank the MGnify service team members – Mahfouz Shehu, Aleksandar Rajkovic, Martin Beracochea and Sandy Rogers, for their technical assistance and code review, and Anna Lopatina for insightful discussions.

## AUTHOR CONTRIBUTIONS

Conceptualization, K.M. and R.D.F.; methodology, K.M., E.K., A.Y., and R.D.F.; Investigation, K.M., A.Y., J.W., and F.C.; writing—original draft, K.M.; writing—review & editing, K.M., F.C., A.Y., A.T., E.K., J.W. A.T. and R.D.F.; funding acquisition, R.D.F.; resources, E.K. and K.M.; supervision, R.D.F.

## DECLARATION OF INTERESTS

The authors declare no competing interests.

## DECLARATION OF GENERATIVE AI AND AI-ASSISTED TECHNOLOGIES

During the preparation of this work, the authors used ChatGPT in order to improve grammar. After using this tool or service, the authors reviewed and edited the content as needed and take full responsibility for the content of the publication.

## SUPPLEMENTAL INFORMATION

**Figures S1–S5 and Table S1-S10**

**Table S1. Comparison of EVADES with dbAPIS and Anti-Defense Finder in terms of sequence diversity and defence families**

**Table S2. Structural similarity between defence proteins**

**Table S3. Uncharacterised ADPs and proteins from counteracted defence systems used in co-folding**

**Table S4. Moderate and high-confidence complexes predicted for uncharacterised ADPs**

**Table S5. Data on protein triggers of bacterial immunity**

**Table S6. DNA mimics**

**Table S7. High-confidence complexes predicted between DNA mimics and defence proteins**

**Table S8. Predicted ipTM/pTM scores for complexes between pairs of NTPs, three ions of Mg^2+^ and YP_009091875 or YP_009031408.**

**Table S9. High-confidence binders of bacterial defence systems from phage hotspots with anti-defence proteins.**

**Table S10. Homologs of ADPs in viral and phage fraction of GenBank and IMG/VR 7.1**

## STAR★METHODS

### KEY RESOURCES TABLE

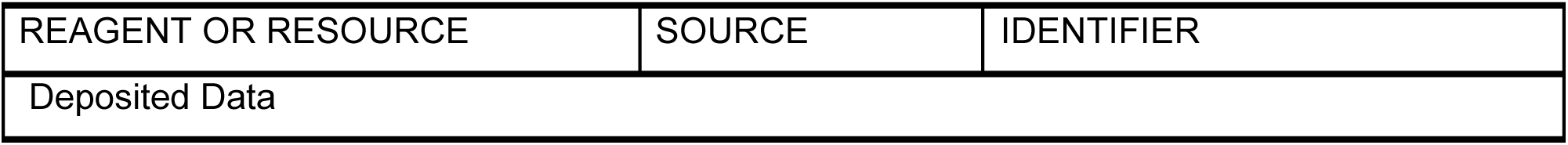
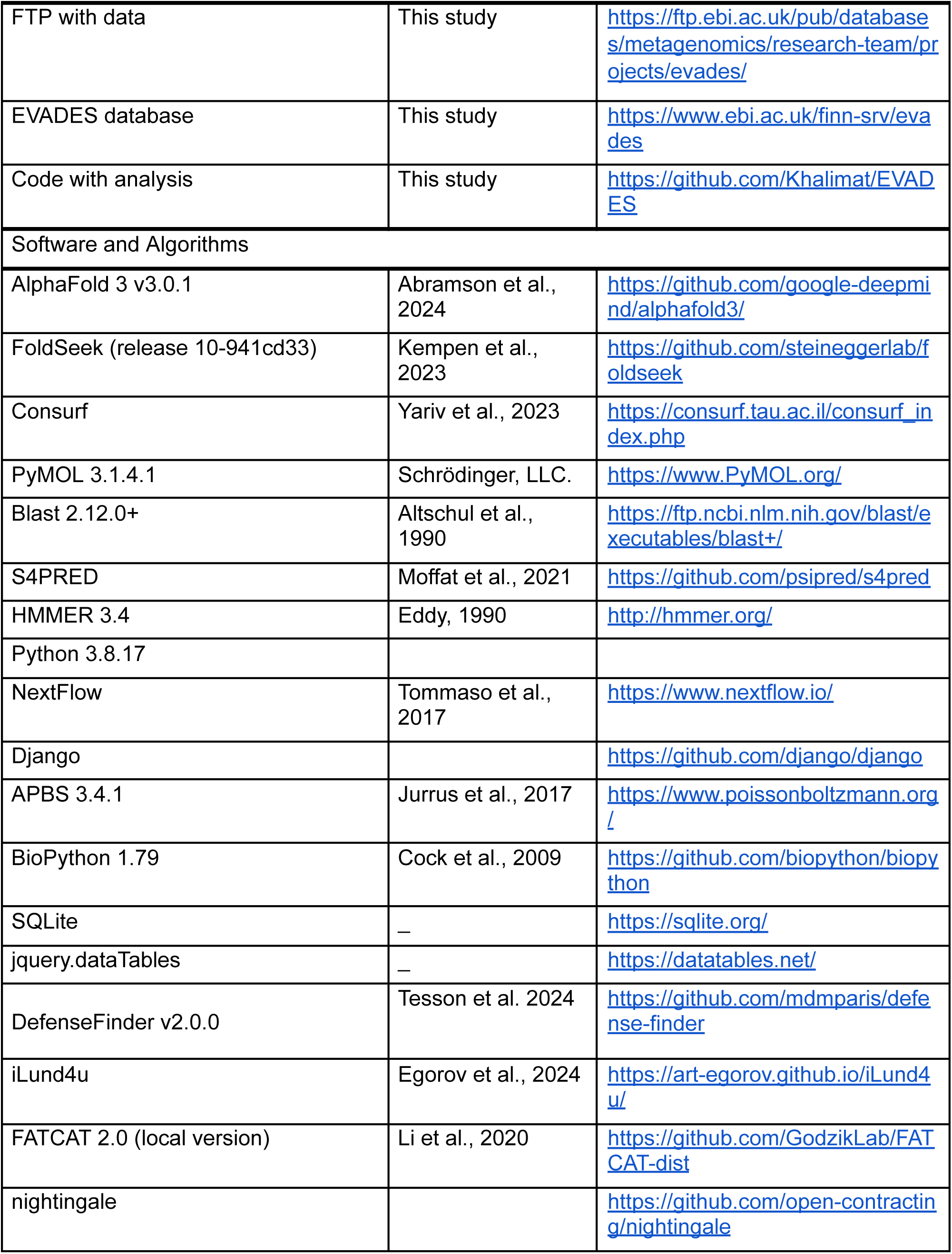
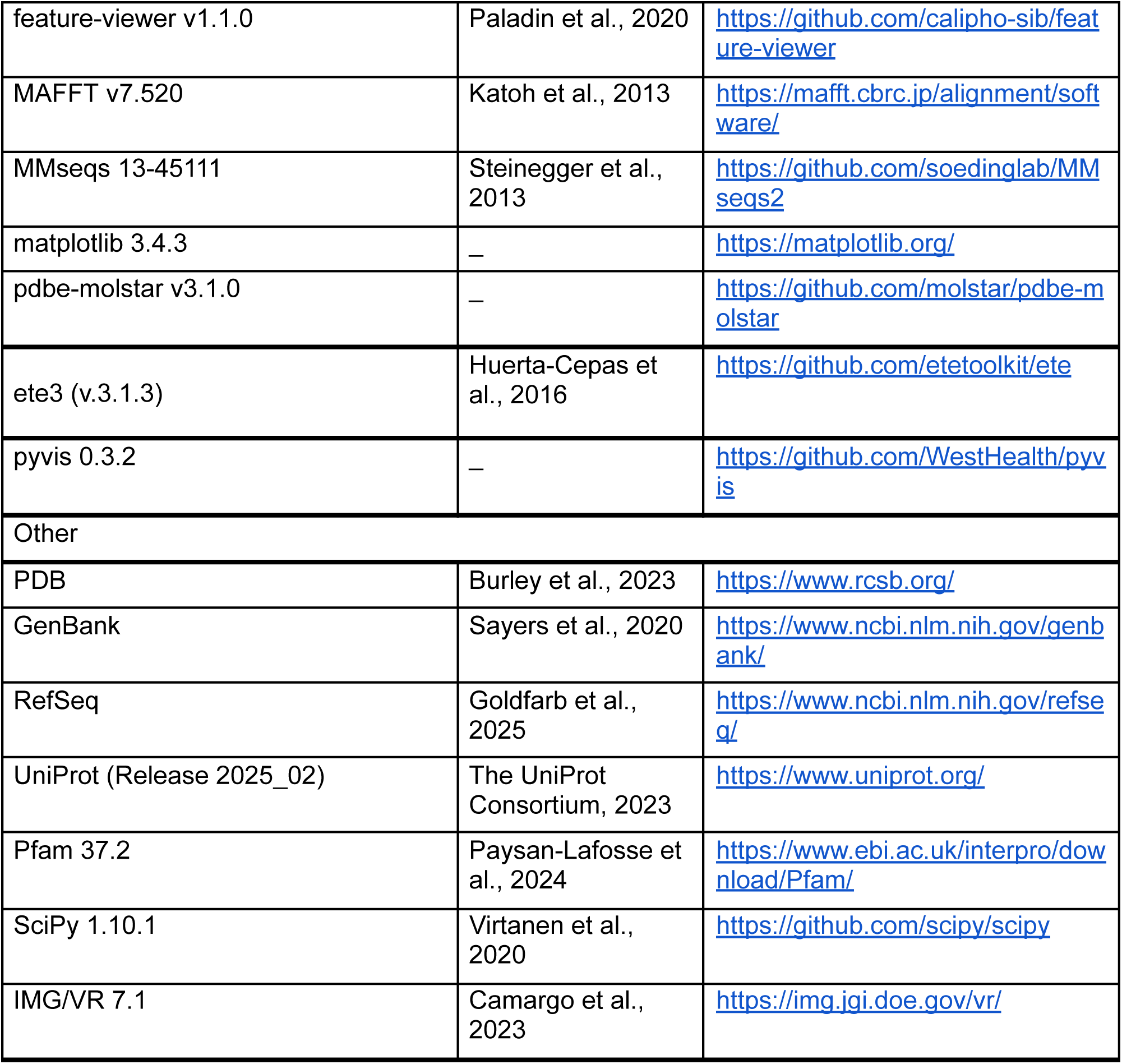

### METHOD DETAILS

#### Data collection

We analysed research literature reporting phage ADPs and extracted information on the defence systems they counteract, MoAs, PDB accessions, data on their potential to trigger other defence systems, and protein sequences. We exclude entries where protein sequences were not provided in the articles, supplementary material or as identifiers in protein or gene databases. For ADPs that function as part of multicomponent systems, we collected and cross-referenced relevant entries accordingly. Additionally, we assigned Evidence and Conclusion Ontology (ECO)^8^ labels to describe evidence supporting entries in our database. We categorised entries in our database based on MoA.

We ran hmmer v3.4^183^ (with option --cut_ga) with Pfam 37.2^184^ to identify existing profile HMMs in Pfam corresponding to ADPs. We produced structural models using AlphaFold 3^38^ for ADPs without a corresponding entry in the PDB. We used AlphaFold 3 v3.0.1 and generated four seeds (1, 2, 3 and 4) through this work. For ADPs without characterised MoA, we searched for structural homolog using FoldSeek^185^ in the pregenerated target databases bundled with FoldSeek (PDB, Alphafold/UniProt and ESMAtlas30, e-value < 0.00001). We used S4PRED to predict protein secondary structure based on the ADP sequences^186^. We used FoldSeek release 10-941cd33 and the following command for the identification of homologous structures through this work: *foldseek easy-search [query_structure] [target_database] [output_file]*.

We downloaded three databases to identify homologs of ADPs: (i) the GenBank database of phage genomes (release from 30-04-2025)^187^, (ii) the GenBank database of viral genomes (release from 30-04-2025), and (iii) IMG/VR version 7.1^188^. We used BLASTp (v2.12.0+)^189^ searches to identify homologs, applying the following criteria to determine significance: e-value < 1^e-5^, query coverage ≥ 80%, and identity ≥ 30%. Additionally, we extracted available phylogenetic information about genomes encoding homologs of ADPs using python package ete3 (v.3.1.3) and from metadata provided in IMG/VR.

We combined the hits obtained from these three databases, added the initial query sequence, de-replicated the sequences using CD-HIT version 4.8.1 at 90% identity and 90% coverage, aligned them using MAFFT version 7.520, manually trimmed the MSAs using Belvu version 4.44.1, and created profile HMMs using *hmmbuild* from the HMMER package. The resulting profile HMM library is available via FTP.

#### Comparison with other resources

To compare ADP coverage in EVADES, dbAPIS^148^, and Anti-Defense Finder^147^, we combined all sequences from these databases, dereplicated them using MMseqs2 (v13.45111, --cov-mode 0)^190,191^ at 90% identity and 90% coverage, and assessed the presence of sequences or closely related homologs (defined as sequences sharing 90% identity over 90% of the query length) in all three databases. Additionally, we calculated the number of ADPs per bacterial defense family and aggregated the results to assess how well EVADES, dbAPIS, and Anti-Defense Finder represent the diversity of ADPs.

#### Creation of the web resource

The web resource was developed using the Python Django framework, and the presented data and metadata are stored in an SQLite database. We created a Nextflow^192^ pipeline to automate SQLite database generation from a json-file with the database. For the visualisations, the following JavaScript libraries were utilised: pdbe-molstar v3.1.0 for the 3D structure viewer, feature-viewer@v1.1.0 for the predicted secondary structure viewer, jQuery DataTables for Pfam annotations, and the Nightingale library^193^ for the interactive Pfam annotations viewer.

#### Sequence similarity comparison

We generated a BLASTp database from all ADP sequences, we conducted homology searches using individual ADP sequences as queries and an e-value threshold of 0.001, and filtered out self-matches. All sequence pairs which passed the e-value threshold, except for two (Lidtsur 6 - Forsur 7 and AcrIIA1 - AcrIB5), exceeded the empirical threshold of 25% identity used to establish homology. To further investigate these exceptions, we used FATCAT 2.0 to assess structural similarity. We used the local version of FATCAT 2.0 and the following command for all pairwise structure similarity comparisons throughout this work: *FATCAT -p1 [pdb_file_1] -p2 [pdb_file_1] -o [pdb_file_1-pdb_file_2] -m -ac -t.* The resulting p-values were: 4.65^e-12^ for Lidtsur 6 -Forsur 7 and 4.38^e-07^ for AcrIIA1 - AcrIB5, so we included them as pairs sharing homology.

#### Similarity with the defence triggers

We compared ADP sequences from our collection with the triggers of bacterial defence systems^28^ using BLASTp (e-value < 0.001)^194^ and analysed the results in order to find new connections to ADPs. We identified multiple matches between the mga47 protein, which inhibits DarTG defence, and several putative Borvo triggers, showing strong sequence similarity. The highest-scoring match exhibited 100% identity across the full length of both sequences. We also observed similarities between promiscuous kinases from phages T7 and JSS1 and putative ShosTA triggers, with the T7 kinase being identical to one of the putative ShosTA triggers. Then we explored the Avs type I trigger (NCBI GenPept accession: WP_317906625.1) from *Bacillus* phage SBSphiC, as it annotated in the original paper as dUMP hydroxymethylase and was not reported as an ADP, but we speculated it may function as a protein involved in DNA hypermodification. We predicted its 3D structure using AlphaFold 3 using the same parameters as previously specified^38^, and searched for homologs in PDB^174^ using FoldSeek^185^. The closest homolog identified was CMP hydroxymethylase MilA (PDB accession: 5JNH). We aligned the structure of MilA with the Avs type I trigger using the local version of FATCAT 2.0 (p-value 0.00^e+00^). Residues K133 and A176 define MilA’s substrate specificity; it has been shown that the mutations K133R and A176S change substrate specificity from CMP to dCMP^156^. We analysed the alignment between the Avs type I trigger and CMP hydroxymethylase MilA in PyMOL (version 3.1.4.1, used throughout this work), identifying the closest residues in the Avs type I trigger that align with those determining substrate specificity in MilA. It has previously *Bacillus* phage SP10, which is closely related to SBSphiC, encodes a dUMP hydroxymethylase involved in DNA hypermodification^195^ (NCBI GenPept accession: YP_007003445.1), which shares 97% sequence identity with the Avs type I trigger. This suggested that dUMP may also serve as a substrate of the type I Avs trigger. To explore this, we co-folded the Avs type I trigger with dUMP, dCMP, and CMP using AlphaFold 3^38^, as ipTM scores for complexes can be indicative of the substrate preference.

We also compared the structures in EVADES to triggers of bacterial defence systems reported by Nagy *et al.* ^155^ using Foldseek. We created query and target databases with the command *foldseek createdb*, performed the comparison using *foldseek search*, converted the output to a TSV file with *foldseek convertalis,* and filtered the results using an e-value threshold of 0.01. However, we did not identify any matches apart from Gam protein, which was previously reported as a trigger and.

We calculated the number of homologs per protein for ADPs known to trigger bacterial defense systems and for non-triggers in the viral and phage fractions of GenBank and IMG/VR (see the Data Collection section). We then compared the two samples using the Mann–Whitney U test from the SciPy package (version 1.10.1)^196^ and calculated the means using the *statistics.mean* function in Python.

#### Prediction of ADPs inhibiting multiple defence systems based on protein sharing network for defence systems

We compiled information from research articles on defence proteins known to interact directly with ADPs. Although direct interactions between anti-Septu proteins (Anhur and Svarog^18^) and the Septu proteins have not been reported, we included PtuA and PtuB in our analysis due to their presence and conservation between Septu defence and type 1A retrons (Ec78, Ec83, Vc95)^197^. Some ADPs with uncharacterised MoA inhibit defence systems composed of a single protein, suggesting that those defence proteins are likely to be targeted—directly or indirectly—by ADPs. Therefore, we also included these defence proteins in our analysis. A full list of the defence proteins used in this experiment is provided in Supplementary Table 2.

We retrieved the structures of these defence proteins from DefenseFinder^198^ (monomer structures) and the PDB, or predicted them if not available in either source. All monomer structures available in DefenseFinder^198^ were included and supplemented with structures of defence proteins from the previous experiment (see Supplementary Table 2) to build a reference database of defence proteins. We used the command *foldseek createdb*^185^ to construct the target database. Structural homologues of ADP-binding proteins were identified using *foldseek easy-search*^185^, applying a stringent e-value cut-off (<1^e-8^) to prioritise functionally relevant structural matches. We then merged the resulting tables, removed redundancy (e.g. multiple entries for the same protein across different defence system subtypes), and visualised the results using the Pyvis 0.3.2 library in Python.

To validate selected structural matches, we used AlphaFold 3 to predict complex formation, with a particular focus on interactions between PglX homologs and Ocr. First, we predicted the structure of the PglX-Ocr complex and assessed whether the ipTM score exceeded the recommended threshold of 0.6. We also predicted interactions between Ocr and four other defence proteins: gp28 (Panchino), drmMII (DISARM), EcoprrI (RM), and HsdM (RM). While predictions for EcoprrI (ipTM = 0.42, pTM = 0.75) and HsdM (ipTM = 0.3, pTM = 0.72) did not meet the threshold, AlphaFold 3 produced moderate-confidence predictions for the Ocr–gp28 complex (ipTM = 0.79, pTM = 0.79, pDockQ = 0.56) and the Ocr–drmMII complex (ipTM = 0.72, pTM = 0.87, pDockQ = 0.401). To calculate the pDockQ score, we implemented a custom script capable of processing cif-files generated by AlphaFold 3. The pDockQ metric was proposed by Bryant *et al.*^199^ as a measure of protein-protein complex prediction quality, with a recommended threshold of pDockQ ≥ 0.23. We visualised both predicted complexes in PyMOL, highlighting residues in gp28 and drmMII that interact with Ocr, defined as residues within 8 Å of any Ocr residue.

#### Prediction of direct interactions between ADPs with uncharacterised mechanisms of action and corresponding defence proteins

First, we obtained the sequences of the defence systems which were inhibited by uncharacterised proteins. We downloaded all monomer PDB-files provided by the DefenseFinder database^198,200^, converted them to fasta-format using BioPython 1.79^201^, and removed duplicate entries. As sequences for CRISPR and RM proteins were absent from the DefenseFinder database, we searched for them in RefSeq^202^, CRISPRCasFinder^203^, UniProt^204^ and PDB^152^. We looked for defence protein sequences with taxonomic annotations corresponding to species specified in the original papers reporting ADPs. We excluded the O-antigen-based barrier, as the polysaccharide chains are highly variable and cannot be reliably modelled. We also excluded ADPs with uncharacterised MoA shown to act indirectly—such as ArdB and KlcA—or are hypothesised to act as negative regulators of anti-defence systems (e.g. ArdK, ArdR, and DdrB). The complete table of sequences for co-folding experiments is provided in Supplementary Table 3. Secondly, we run the AlphaFold 3^38^ v3.0.1 data pipeline (*--run_inference=false*) on individual defence and anti-defence protein sequences to generate MSA and structural templates. This approach allowed reuse of MSA/template data for proteins appearing multiple times. Third, we created JSON files combining the MSAs and templates from the previous step for each corresponding defence and anti-defence protein pair. We then ran AlphaFold 3^38^ inference with 4 seeds (1, 2, 3, 4), producing 20 models and focused only on those complexes where both the average ipTM and the best model’s ipTM were greater than 0.6. We then computed the pDockQ scores for the best models. Predicted complexes were classified into high-confidence (best model ipTM ≥ 0.8) and moderate-confidence (0.6 < best model ipTM < 0.8). All results are available in Supplementary Table 4.

#### Putative MoA for AcrIC5

We predicted a high-confidence (ipTM = 0.92, pTM = 0.85, pDockQ = 0.56) complex between AcrIC5 and *Pseudomonas aeruginosa* Cas8c (*Pa*Cas8c, NCBI GenPept accession: UEM35121.1). To gain mechanistic insights from the predicted complex between AcrIC5 and *Pa*Cas8c, we sought to annotate the region in Cas8c that binds AcrIC5. We looked for the closest homolog in the PDB using FoldSeek^185^ and found a Cascade complex containing a Cas8c homolog from *Desulfovibrio vulgaris* (PDB accession: 7KHA)^153^. We aligned it using FATCAT 2.0 with the structure of *Pa*Cas8c predicted with AlphaFold 3^38^ and found that the binding site for AcrIC5 is extensive and includes the putative PAM-binding site (residues: GVDAKGK). We visualised the results in PyMOL, colouring AcrIC5 based on electrostatic potential (using Adaptive Poisson-Boltzmann Solver, APBS, Electrostatics module^205^).

#### Putative MoA for AcrIC3

We predicted a high-confidence complex (ipTM = 0.91, pTM = 0.90, pDockQ = 0.633) between AcrIC3 and *Pseudomonas aeruginosa* Cas3 (*Pa*Cas3, NCBI GenPept: UEM35119.1). To annotate *Pa*Cas3 protein, we took its structure predicted by AlphaFold 3^38^ and looked for structural homologs in PDB using FoldSeek^185^. We identified Cas3 from *Thermobifida fusca* YX (PDB accession: 4QQZ) as a closest homolog (e-value = 1.74^e-18^)^206^. Then we aligned the structures using FATCAT 2.0, and relying on the domain coordinates reported for *Thermobifida fusca’*s Cas, annotated domains (1-216 HD, 217-430 RecA1, 431-602 RecA2, 603-649 Linker, and 650-744 CTD) in *Pa*Cas3. We visualised the complex between AcrIC3 and *Pa*Cas3 in PyMOL. We used the BioPython 1.79^201^ module NeighborSearch to identify residues in Cas3 likely interacting with residues in AcrIC3 (defined as residues in 8Å or less). We used matplotlib (v.3.4.3) to visualise the domain architecture, plotting grey dots on residues in *Pa*Cas3 interacting with AcrIC3.

#### Putative MoA for AcrIB4

We predicted a high-confidence complex (ipTM = 0.91, pTM = 0.76, pDockQ = 0.424) between AcrIB4 and *Listeria seeligeri* Cas8 (*Ls*Cas8b1) with its N-terminal residues (504–530) forming the binding interface. To annotate the *Ls*Cas8b1 binding interface, we took its AlphaFold 3^38^ predicted structure and looked for structural homologs in the PDB using FoldSeek^185^. However, as we did not identify any matches, we manually looked for type I-B Cascade complexes containing Cas8 in the PDB and found a complex (PDB accession: 8FCJ) where Cas8 protein shares high similarity to *Ls*Cas8b1^207^. We found that the Cas8 residues corresponding to the AcrIB4 binding interface in *Ls*Cas8b1 are involved in binding Cas11 in the type I-B Cascade. We visualised the complexes between AcrIB4-*Ls*Cas8b1 and Cas8-Cas11 in PyMOL.

#### Putative MoA for AcrIIA26

We predicted a high-confidence complex (ipTM = 0.87, pTM = 0.79, pDockQ = 0.634) between AcrIIA26 and *Spy*Cas9 (NCBI GenPept accession: QHB64844.1). We wanted to explore the positioning of AcrIIA26 relative to sgRNA and target DNA, so we took sgRNA and target DNA sequences from a PDB entry with accession 4OO8^208^, co-folded them with *Spy*Cas9 using AlphaFold 3^38^ and predicted a very high-confidence complex (ipTM = 0.94, pTM = 0.92). We positioned the AcrIIA26-*Spy*Cas9 complex and the complex between sgRNA, target DNA and *Spy*Cas9 similarly in space using PyMOL, which allowed us to infer that AcrIIA26 occludes the DNA binding site. We calculated electrostatic potential on AcrIIA26 using APBS Electrostatics module in PyMOL and visualised its surface interacting with *Spy*Cas9.

#### Structure based search for anti-defence proteins in eukaryotic viral protein database

We conducted structure-based homology search for phage ADPs in a eukaryotic viral protein database^160^ using FoldSeek^185^ (considering the size of DB, 68K entries, we filtered on e-value < 0.00001). Then, we analysed the results and focused on matches with ADPs counteracting defence systems shared between bacteria and eukaryotes, namely anti-Avs proteins ORF83 and ORF55^17^ and NTase^53^ inhibiting TIR and STING immune response.

#### ORF55 functional annotation and homologs in eukaryotic viruses

We looked for homologs of ORF55 using its structural model in PDB^152^ with FoldSeek^185^ and found that the closest match is the DNA ligase (PDB accession: 2Q2U) from *Chlorella* virus (eukaryotic DNA viruses). We superimposed ORF55 and the DNA ligase from *Chlorella* virus using FATCAT 2.0 (p-value = 0.00^e+00^) to obtain a more accurate estimate of similarity. We analysed the paper reporting the structure of the DNA ligase from the *Chlorella* virus and retrieved active site residues responsible for its catalytic activity. Then we used the superimposed complex of ORF55 with the *Chlorella* virus DNA ligase obtained using FATCAT, analysed the alignment in PyMOL, and retrieved the residues in ORF55 aligning with the active site residues in the *Chlorella* virus DNA ligase.

We modelled interaction between ORF55 and the DNA fragment from the *Chlorella* virus DNA ligase using AlphaFold 3^38^ and predicted a high confidence complex. Next we used the same DNA fragment, but introduced a nick, and one- and two-nucleotide gapes and co-folded with ORF55 and ATP, where similar to the *Chlorella* virus DNA ligase, ORF55 showed preference for nicked (ipTM = 0.95, pTM = 0.95) and one-nucleotide gapped DNA (ipTM = 0.95, pTM = 0.95) compared to two-nucleotide gapped DNA (ipTM = 0.87, pTM = 0.91).

#### ORF83 functional annotation and homologs in eukaryotic viruses

We looked for structural homologs of ORF83 in the PDB database using FoldSeek^185^, aligned the top four matches using FATCAT 2.0 and identified *Cg*1261 (PDB ID: 5WQ3, 4.39^e-7^) – the type II cytokine activating protein from LOG family (The Lonely Guy) as the highest scoring match. We analysed the paper with functional characterisation of *Cg*1261 and found it was reported to hydrolyze AMP into an adenine base and ribose 5-phosphate. So, we modelled interaction between ORF83 and AMP using AlphaFold 3^38^. Then we aligned the ORF83-AMP complex with *Cg*1261 using FATCAT 2.0, selected active site residues in *Cg*1261 from the paper^209^ and found closest aligning residues in ORF83. Then we predicted the structure of ORF83’s homolog from eukaryotic dsDNA viruses (NCBI GenPept accession: YP_009506880) in complex with AMP, aligned it with ORF83 using FATCAT 2.0, and visualised the results in PyMOL.

#### NTase

We aligned the structural model of *Bacillus* phage Bcp1 NTase with its homolog we identified in *Erinnyis ello* granulovirus (NCBI GenPept accession: YP_009091875.1) using FATCAT 2.0 (p-value = 2.74^e-10^)^210^ and visualised the results in PyMOL. Next, we calculated the number of positions in the phage NTase and its homolog from *Erinnyis ello* granulovirus matching the catalytic motif of phage NTases and visualised the results.

We aimed to profile substrate specificity of phage and viral NTases, so we modelled^38^ the complexes between one of the NTases and pairs of NTPs (we only used ATP, GTP, or UTP, as they are typically utilised by CD-NTases) and 3 ions of Mg^2+^, which have been reported as cofactors stabilising donor and acceptor NTPs^177^. We visualised the ipTM and pTM scores as heatmaps. We used PyMOL to visualise the hydrogen bonds formed between *Erinnyis ello* granulovirus NTase, two molecules of UTP and three ions of Mg^2+^.

#### Prediction of putative ADPs from diversity hotspots

Diversity hotspots are genomic regions characterised by elevated variability, typically flanked by conserved genes, and frequently observed in phage and plasmid genomes. These regions arise through mechanisms such as mobile genetic element insertions, recombination, and hypermutation, and often carry accessory genes involved in defence, anti-defence, virulence, and antimicrobial resistance^179^. We ran iLund4u to identify ADPs encoded such diversity hotspots in phages (*iLund4u protein -fa [adp].fa -db phages -o [adp]_phages*) using the default parameters. iLund4u uses a tool named Pharokka^211^ to annotate proteins from diversity hotspots replying on profile HMMs derived from the database Prokaryotic Virus Remote Homologous Groups (PHROG)^212^. We selected all uncharacterised proteins based on iLund4u results from phage diversity hotspots that encode ADPs in this analysis. As the iLund4u results also contain information about cluster membership for proteins, we selected protein clusters containing more than 50 members. These clusters were further refined by re-clustering the proteins at 40% sequence identity over 80% of their length. To exclude proteins with other known functional annotations, we used HMMER with Pfam version 37.2 and the *--cut_ga* option on cluster representatives. These steps enabled us to focus on uncharacterised proteins from diversity hotspots for structural modelling. We first modelled individual phage proteins using a local version of AlphaFold 3, generating four models with seeds 1, 2, 3, and 4. Proteins with an average predicted local Distance Difference Test (pLDDT) score below 70 were filtered out.

Next, we performed a three-step co-folding process to identify potential interactions between phage proteins and bacterial defence proteins.We aimed to select pairs of proteins that consistently showed high confidence scores across different seed numbers, so we randomly generated a different seed number for each step:

1. Initial Co-folding: We co-folded all possible phage–bacterial protein pairs (255 × 608 = 155,040 combinations) using local AlphaFold 3 with seed 203490234, generating five models per pair. We retained pairs where at least three of the five models had a predicted confidence score (0.8 × ipTM + 0.2 × pTM) above 0.8.
2. Server-based Co-folding: The shortlisted pairs were co-folded again using the AlphaFold 3 server (between 27 May and 20 June 2025) with seed 349024. We applied the same filtering criterion: a confidence score above 0.8 in at least three out of five models.
3. Final Co-folding with Multiple Seeds: Pairs passing the second step were modelled again locally with AlphaFold 3 using five different seeds (3294, 938934343, 778343, 343, and 53343). We selected those with an average confidence score above 0.8 in at least 17 out of the 25 total models.

Finally, for each selected protein pair, we calculated the pDockQ score and retained only those with a pDockQ score ≥ 0.23.

### QUANTIFICATION AND STATISTICAL ANALYSIS

The p-values and e-values produced by tools are described in the text with references to original articles.

We used the Mann-Whitney U-test to compare the distributions in Figure 3B, and the Fisher exact test to calculate the Odds ratio and p-value in Figure 3C.

We calculated correlation between ipTM scores for phage and viral NTase using Pearson correlation coefficient.

## Notes

https://www.ebi.ac.uk/finn-srv/evades

### Competing Interest Statement

The authors have declared no competing interest.

### Summary of Updates

Mostly grammar changes and supplemental files updated.

